# Investigating a Relation between Amyloid Beta Plaque Burden and Accumulated Neurotoxicity Caused by Amyloid Beta Oligomers

**DOI:** 10.64898/2026.04.07.717091

**Authors:** Andrey V. Kuznetsov

## Abstract

Alzheimer’s disease (AD) is characterized by the accumulation of amyloid-β (Aβ), yet the specific link between plaque burden and cognitive decline remains a subject of intense investigation. This paper presents a mathematical model that simulates the coupled dynamics of Aβ monomers, soluble oligomers, and fibrillar species in the brain tissue. By modifying existing moment equations to include a dedicated conservation equation for Aβ monomers, the model explores how various microscopic processes, such as primary nucleation, surface-catalyzed secondary nucleation, fibril elongation, and fragmentation, contribute to macroscopic disease progression. Central to this study is the concept of “accumulated neurotoxicity” as a surrogate marker of biological age, defined as the time-integrated concentration of soluble Aβ oligomers. Unlike plaque burden, accumulated neurotoxicity cannot be reversed, and the harm it causes depends critically on the sequence of events that produced it. Numerical results demonstrate that while plaque burden and neurotoxicity both increase over time, their relationship is non-linear and highly sensitive to the efficiency of protein degradation machinery. Specifically, impaired degradation leads to a rapid advancement of biological age relative to calendar age. The model further identifies oligomer dissociation and fibril fragmentation as potential protective mechanisms that can counterintuitively reduce neurotoxic burden by diverting monomers away from the soluble oligomer pool. These findings provide a quantitative framework for understanding why individuals with similar plaque burdens may experience vastly different cognitive outcomes, underscoring the importance of targeting soluble oligomers early in therapeutic interventions.

## 1. Introduction

Despite extensive evidence of Aβ accumulation in Alzheimer’s disease, its quantitative relationship to cognitive decline remains unresolved. The failure of numerous clinical trials targeting brain Aβ reduction suggests that, to be effective, anti-Aβ therapies must be initiated well before the onset of clinical symptoms, and that achieving this requires a thorough understanding of the dynamics of Aβ deposition [1]. Mounting evidence suggests that soluble Aβ oligomers, rather than insoluble plaques, are the primary drivers of neuronal damage. The clinical trial record indicates that only agents targeting soluble Aβ oligomers have shown clinical efficacy, while those predominantly targeting monomers or plaques have failed to demonstrate a meaningful benefit [2]. Consistent with this, ref. [3] found that the ratio of Aβ oligomer levels to plaque density fully distinguished demented from non-demented patients, with no overlap between groups, demonstrating that two individuals with equivalent plaque burdens can have markedly different cognitive outcomes depending on their respective levels of soluble oligomers. These observations motivate the development of mathematical models that can describe the coupled dynamics of Aβ monomers, oligomers, and fibrils, and thereby provide a quantitative framework for understanding disease progression and informing the design of therapeutic interventions.

The Aβ master equation was formulated in ref. [4] as a nonlinear rate law integrating microscopic processes, including primary nucleation, fibril elongation, and surface-catalyzed secondary nucleation, to link molecular mechanisms with macroscopic aggregation kinetics. The corresponding moment equations were derived in ref. [4] by summing this master equation over aggregate size, reducing the full population balance to equations for the total number and mass concentrations of aggregates. Ref. [5] extended this framework by generalizing the moment equations of ref. [4] to explicitly account for non-fibrillar oligomeric intermediates and the multiple reaction pathways they undergo, including structural conversion, dissociation, and surface-catalyzed association. The moment equations of ref. [5] differ from the Finke-Watzky model [6,7] in that they explicitly incorporate multiple reaction pathways for non-fibrillar oligomeric intermediates, such as dissociation back to monomers and secondary association catalyzed by fibril surfaces. Furthermore, in contrast to the models proposed in refs. [8-10], the equations of ref. [5] explicitly account for fibril length heterogeneity, elongation, and fragmentation, as well as the conversion of oligomers into fibrils via monomer addition.

This paper presents a mathematical model of Aβ oligomer and fibril formation based on a modification of the equations developed in ref. [5]. The principal modification consists of deriving an additional conservation equation for the Aβ monomer species.

## 2. Materials and models

### 2.1. Mathematical framework for Aβ aggregation

The conservation of Aβ monomers, oligomers, fibrillar species of all lengths, and total fibril mass is formulated within a cubic control volume (CV) that receives monomers through one face, while the opposite face is treated as impermeable (Fig. 1). The side length of the CV is *L* ≈ 50 μm, corresponding to approximately half the average distance between Aβ plaques. Applying conservation of free Aβ monomers within the CV and normalizing both sides by the CV’s volume, *L*^3^, yields the following governing equation:

**Fig. 1.**
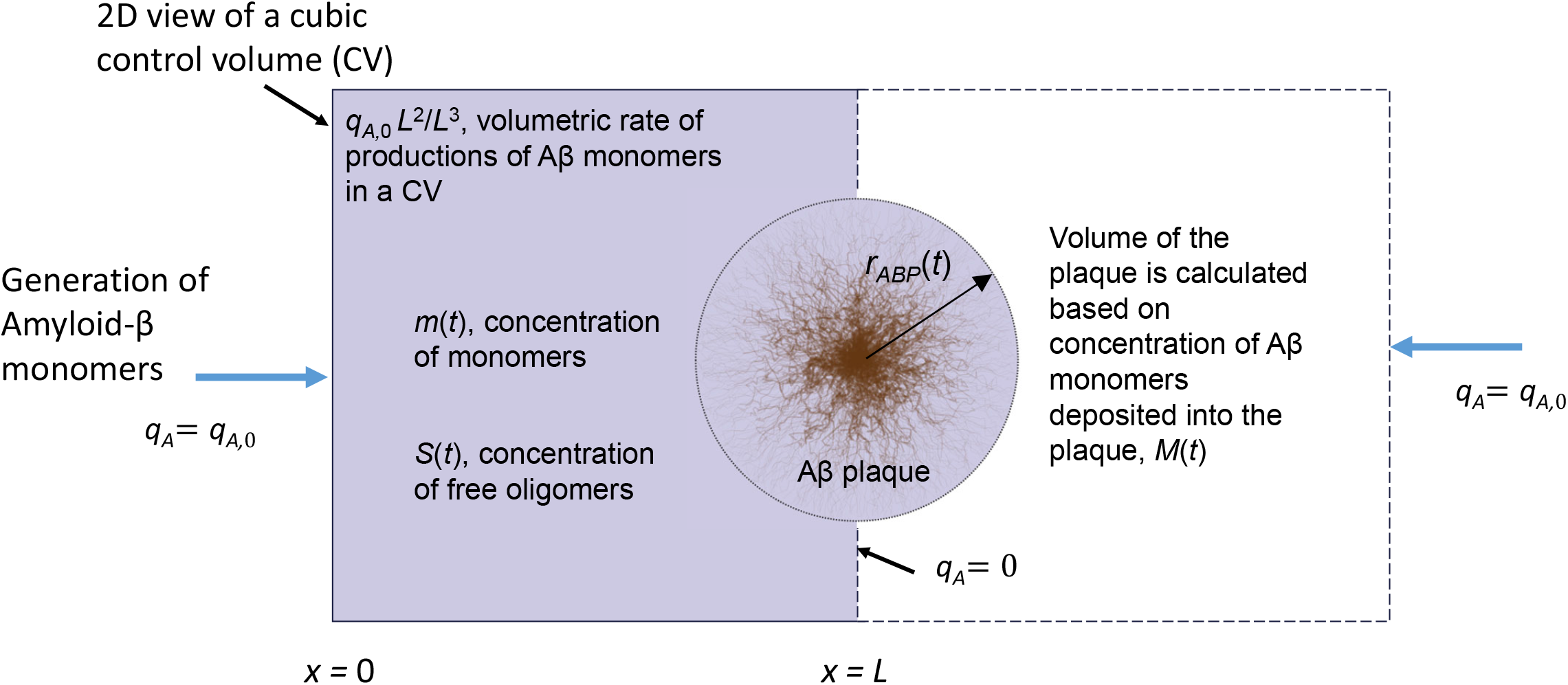
Two-dimensional projection of a cube-shaped control volume (CV) with side length *L*, used to model the aggregation of Aβ monomers into free oligomers and the subsequent formation of elongation-competent fibrillar species of varying lengths. Aβ monomers are produced at the *x* = 0 boundary, while the *x* = *L* boundary is treated as symmetric, admitting no monomer flux. The Aβ plaque is assumed to form at the symmetric interface between two adjacent CVs. A lumped capacitance approximation is adopted, under which all Aβ concentrations are treated as functions of time alone.

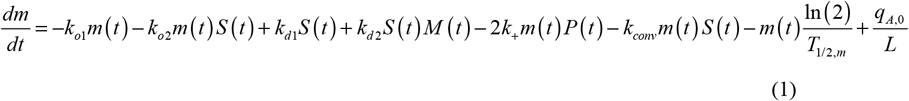

where *t* is the time elapsed since AD onset. The terms on the right-hand side of Eq. (1) each represent a distinct physical process. The first term simulates monomer loss due to nucleation, whereby free monomers are consumed in the formation of oligomers. The second term accounts for monomer loss through secondary, or autocatalytic, conversion: a process in which existing oligomers catalyze the production of new ones. The third term describes monomer gain due to spontaneous dissociation of oligomers into monomers, while the fourth term accounts for monomer gain due to oligomer dissociation catalyzed by fibril surfaces. The fifth term describes monomer depletion arising from fibril elongation: since each fibril possesses two free tips at which monomers can attach, a factor of 2 appears in this term. The sixth term represents the conversion of oligomers into elongation-competent fibrils; the product of *m*(*t*) and *S* (*t*) reflects the fact that this conversion likely proceeds via monomer addition [5]. The seventh term accounts for monomer clearance through proteolytic degradation, where the half-life of the monomers, *T*_1/ 2,*m*_, governs the rate of this process [11]. Finally, the eighth term describes the production of Aβ monomers at lipid membranes, with 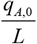 denoting the average volumetric generation rate of monomers within the CV.

Consistent with the Finke-Watzky model, an oligomer is interpreted as an activated monomer and is assumed to carry the same mass as a free monomer [6,7]. This represents a considerable simplification of the oligomer formation mechanism. Applying the conservation principle to the number of free on-pathway Aβ oligomers within the CV and normalizing the resulting equation by the CV’s volume yields:

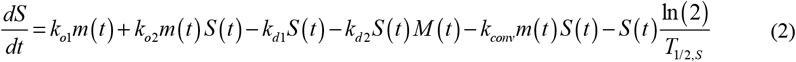

The first two terms on the right-hand side of Eq. (2) represent the gain of Aβ oligomers through nucleation and autocatalytic conversion, respectively. The first two terms are equal in magnitude but opposite in sign to the first two terms on the right-hand side of Eq. (1). The third and fourth terms describe oligomer loss due to spontaneous dissociation into monomers and fibril-catalyzed dissociation, respectively, and are equal in magnitude but opposite in sign to the corresponding terms in Eq. (1). The fifth term describes the loss of oligomers through their conversion into elongation-competent fibrils. The sixth term accounts for oligomer loss due to proteolytic degradation, the rate of which is governed by the oligomer half-life, *T*_1/ 2,*S*_.

Applying the conservation principle to the number of Aβ fibrillar species of all lengths and normalizing by the CV’s volume yields the governing equation for the total fibril concentration, regardless of length:

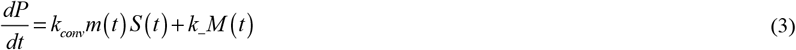

The first term on the right-hand side of Eq. (3) corresponds to the sixth term in Eq. (1) and the fifth term in Eq. (2), but with opposite signs, and represents the conversion of oligomers into elongation-competent fibrils. This process is assumed to proceed via monomer addition, whereby one monomer and one oligomer combine to produce a new fibril. Fibril degradation is neglected in this equation, as it is considered inefficient relative to the rapid clearance of monomeric Aβ [12]. The second term on the right-hand side of Eq. (3) represents the generation of new fibrils through fragmentation of existing ones.

The mass of fibrils within the CV is characterized by the total number of Aβ monomers incorporated into them. Applying conservation of this quantity and normalizing by the CV’s volume yields the following equation:

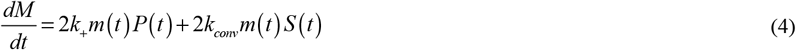

In Eq. (4), the first term on the right-hand side describes fibril elongation through monomer addition, which can occur at either tip of the fibril. The second term describes the conversion of oligomers into fibrils, a process that involves monomer addition to the oligomer. The factor of 2 in this term reflects the incorporation of both a monomer and an oligomer into the fibril mass during this event.

Because the pathophysiological processes of AD unfold over decades [13], the long-term behavior of Aβ concentrations in the CV is effectively independent of initial conditions on this timescale, and all Aβ species can therefore be set to zero at the time of AD onset, *t* = 0. This leads to the following initial conditions for Eqs. (1)-(4):

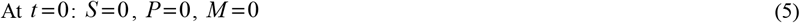

### 2.2. Modelling the growth of plaque volume and radius

The growth model of plaque volume and radius follows ref. [25]. The growth of an Aβ plaque (Fig. 1) is tracked by calculating the total number of Aβ monomers, *N*, incorporated into the plaque over time *t*. Following the approach of ref. [26], this quantity is obtained from the average concentration of Aβ monomers deposited into the plaque:

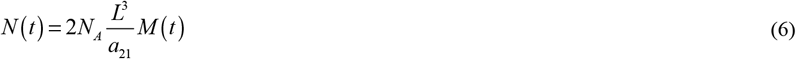

where is *N*_*A*_ Avogadro’s number. The factor of 2 on the right-hand side of Eq. (6) reflects the assumption that the Aβ plaque forms between two adjacent neurons, each of whose membranes releases Aβ monomers at a rate of *q*_*A*,0_, thereby contributing to plaque growth (Fig. 1).

An alternative approach, also following ref. [26], determines *N* (*t*) through the volume of a single inclusion body, *V*_*ABP*_ :

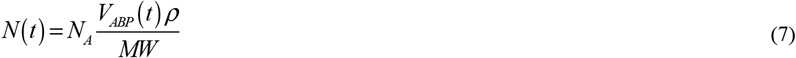

where *MW* denotes the average molecular weight of an Aβ monomer. Equating the right-hand sides of Eqs. (6) and (7) and solving for the plaque volume gives:

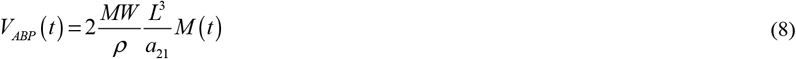

where 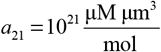 is the conversion factor from 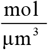 to μM. Assuming the Aβ plaque is spherical,

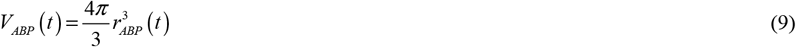

Equating the right-hand sides of Eqs. (8) and (9) and solving for the plaque radius yields:

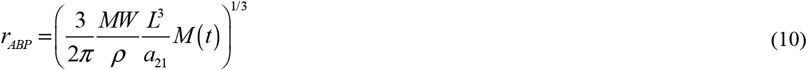

The percentage of the CV occupied by the Aβ plaque is given by:

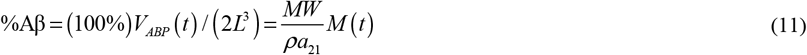

### 2.3. Accumulated neurotoxicity as a surrogate marker of biological age

The neurotoxicity criterion is based on the time-integrated concentration of soluble Aβ oligomers, rather than on instantaneous plaque burden. This choice is motivated by a substantial body of experimental and clinical evidence. Research uncovering the role of soluble Aβ in AD goes back to at least 1999, when ref. [27] demonstrated that the concentration of soluble Aβ is elevated threefold in AD and correlates far more closely with markers of disease severity than does the number of amyloid plaques. Subsequent work has established that Aβ oligomers exert neurotoxic effects through multiple mechanisms, including direct interactions with synaptic receptors that contribute to progressive synaptic failure [28-31], and there is now a large body of literature supporting the view that soluble Aβ oligomers are the primary neurotoxic species in AD [32]. Critically, the damage inflicted by oligomers accumulates over time: it is not the instantaneous oligomer concentration, but its integral over the disease course, that determines the cumulative cellular insult. This notion is supported by ref. [33], who found that the yearly rate of change in Aβ burden improves the prediction of cognitive decline beyond what a single static measurement can provide, underscoring the prognostic value of the temporal trajectory of Aβ accumulation over any snapshot measurement of plaque burden. Accumulated neurotoxicity is therefore defined following ref. [8] as:

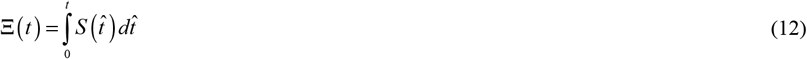

The proposed criterion Ξ can serve as a marker of biological age because its irreversible and path-dependent nature reflects the cumulative, nonlinear progression of neurodegenerative damage. Unlike state-dependent markers that capture only instantaneous protein levels, its irreversibility ensures that even complete clearance of fibrillar deposits cannot reset the “biological clock” to zero, recognizing that past physiological insults leave a lasting imprint on neuronal health. Moreover, its path dependence implies that the criterion Ξ is determined by the history of oligomer exposure rather than solely by the final concentration, thereby capturing both the duration and intensity of toxic stress experienced by the brain. By framing biological age in terms of the aggregation of intrinsically disordered proteins, as proposed by ref. [34], this criterion is consistent with the role of aging as the primary risk factor for AD.

Following ref. [10], the biological age of neurons (hereafter simply referred to as biological age) is defined as:

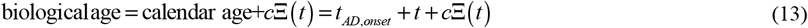

### 2.4. Approximate analytical solution for the limiting case of *k*_*d*1_ = 0, *k*_*d*2_ = 0, *k*_−_ = 0, *T*_1/ 2,*m*_ →∞, and T_1/ 2,*S*_ →∞

Adding Eqs. (1), (2), and (4) for this case yields:

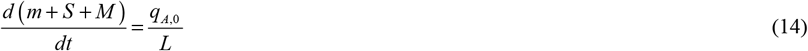

The numerical results indicate that *m* + *S* = *M* (see Figs. 2b, 3b, and 5b). Integrating Eq. (14) yields:

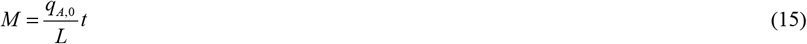

**Fig. 2.**
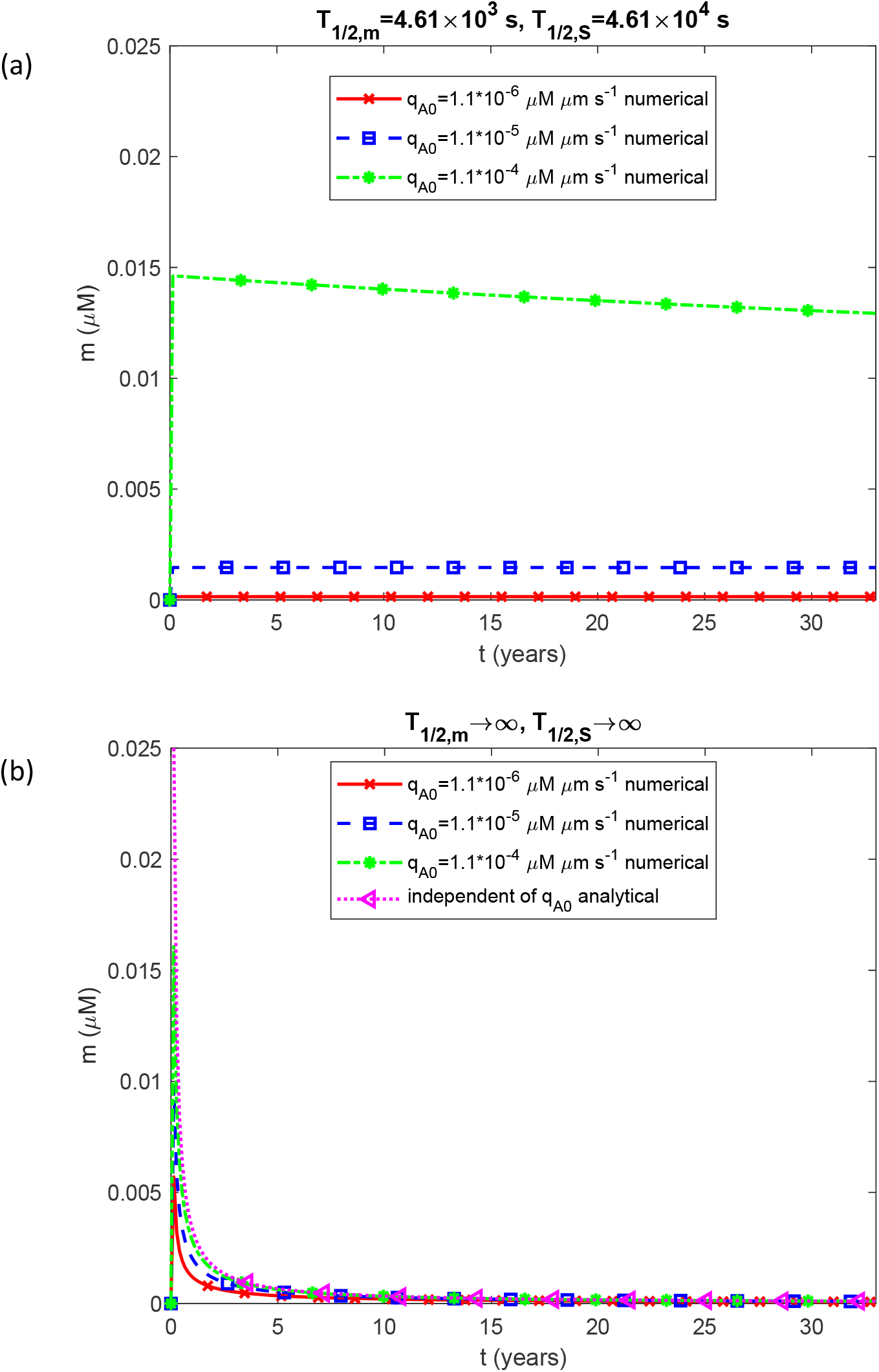
Molar concentration of free Aβ monomers as a function of time, *m*(*t*). The case of negligible oligomer dissociation into monomers (*k*_*d*1_ = 0, *k*_*d*2_ = 0) and negligible fibril fragmentation (*k*_−_ = 0).

The numerical results (see Figs. 2b, 3b, and 4b) suggest that the time dependence of *m, S*, and *P* can be approximated as follows:

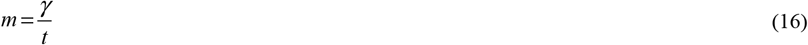

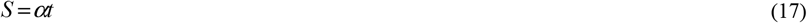

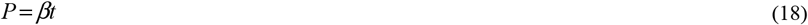

where *α, β*, and *γ* are constants. Substituting this into Eq. (3) yields:

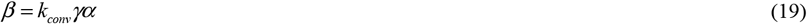

Adding Eqs. (1) and (2) and neglecting the term involving *k*_*conv*_ on account of the small magnitude of *k*_*conv*_ (Table 2) yields:

**Table 1.** Dependent variables used in the model.

| Symbol | Definition | Units |
| --- | --- | --- |
| $m(t)$ | Molar concentration of free Aβ monomers | $\mu\text{M}$ |
| $M(t)$ | Molar concentration of Aβ monomers incorporated into all fibrillar species, regardless of length (a measure of the total fibril mass expressed in monomer units) | $\mu\text{M}$ |
| $P(t)$ | Molar concentration of all fibrillar species, regardless of length (a measure of the total number of fibrils) | $\mu\text{M}$ |
| $S(t)$ | Molar concentration of free on-pathway Aβ oligomers | $\mu\text{M}$ |

**Table 2.**
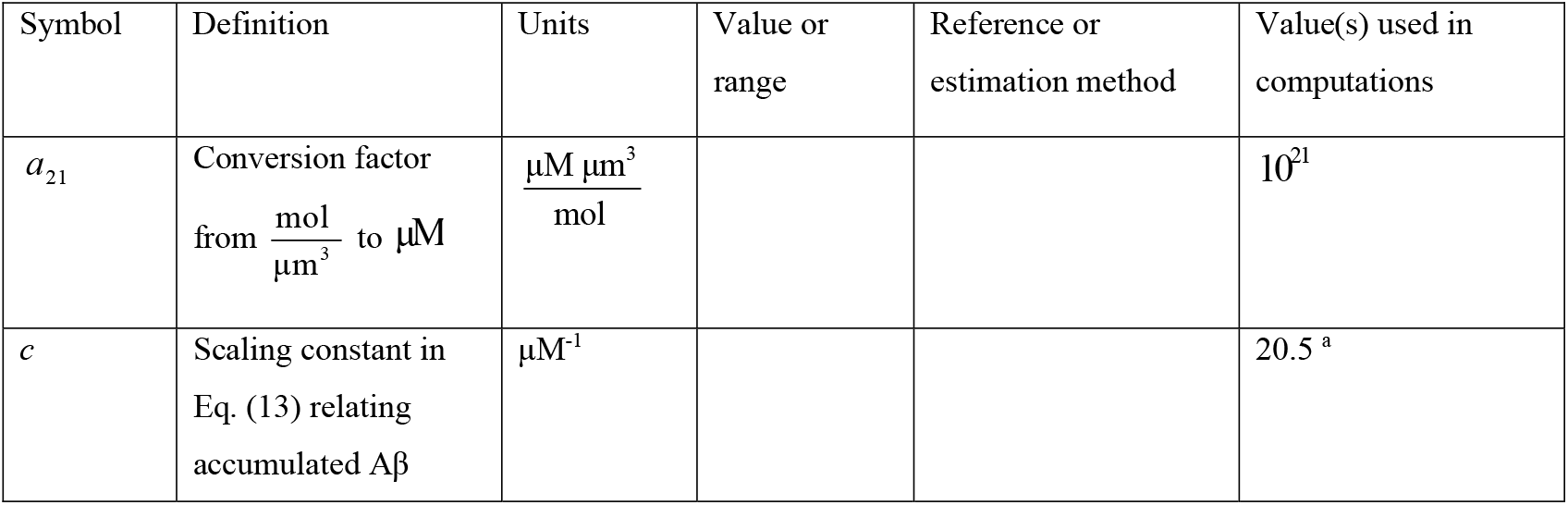
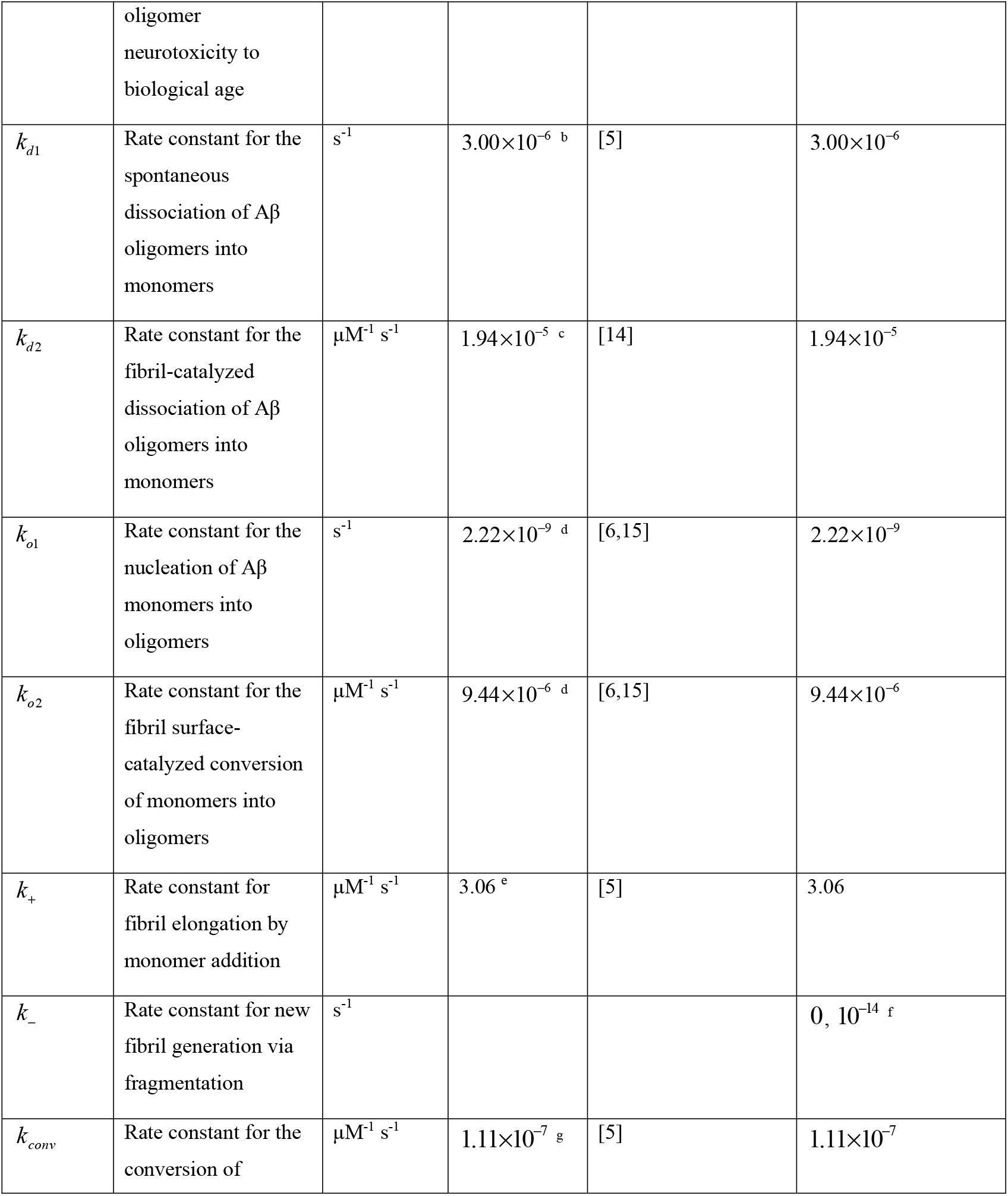
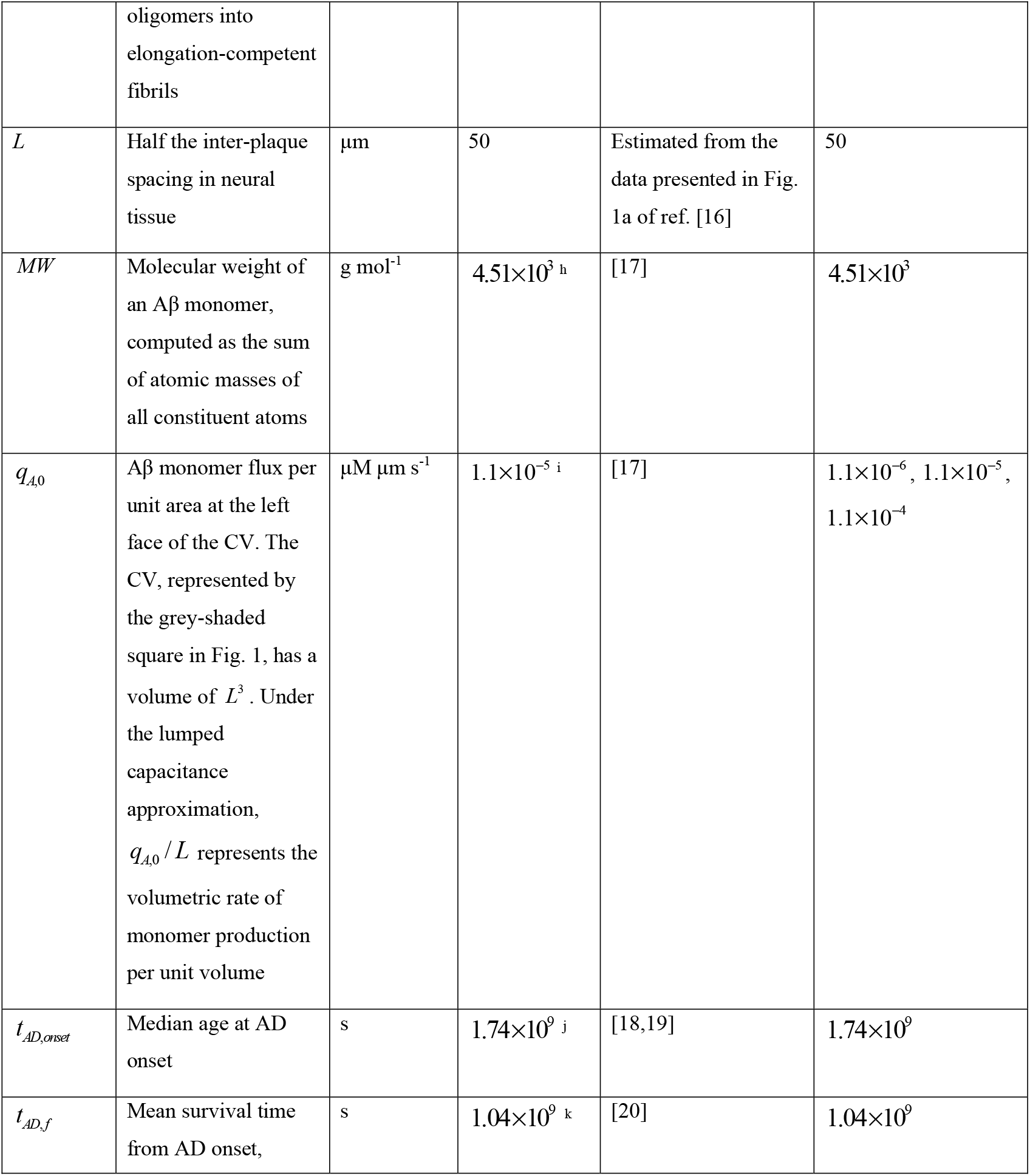
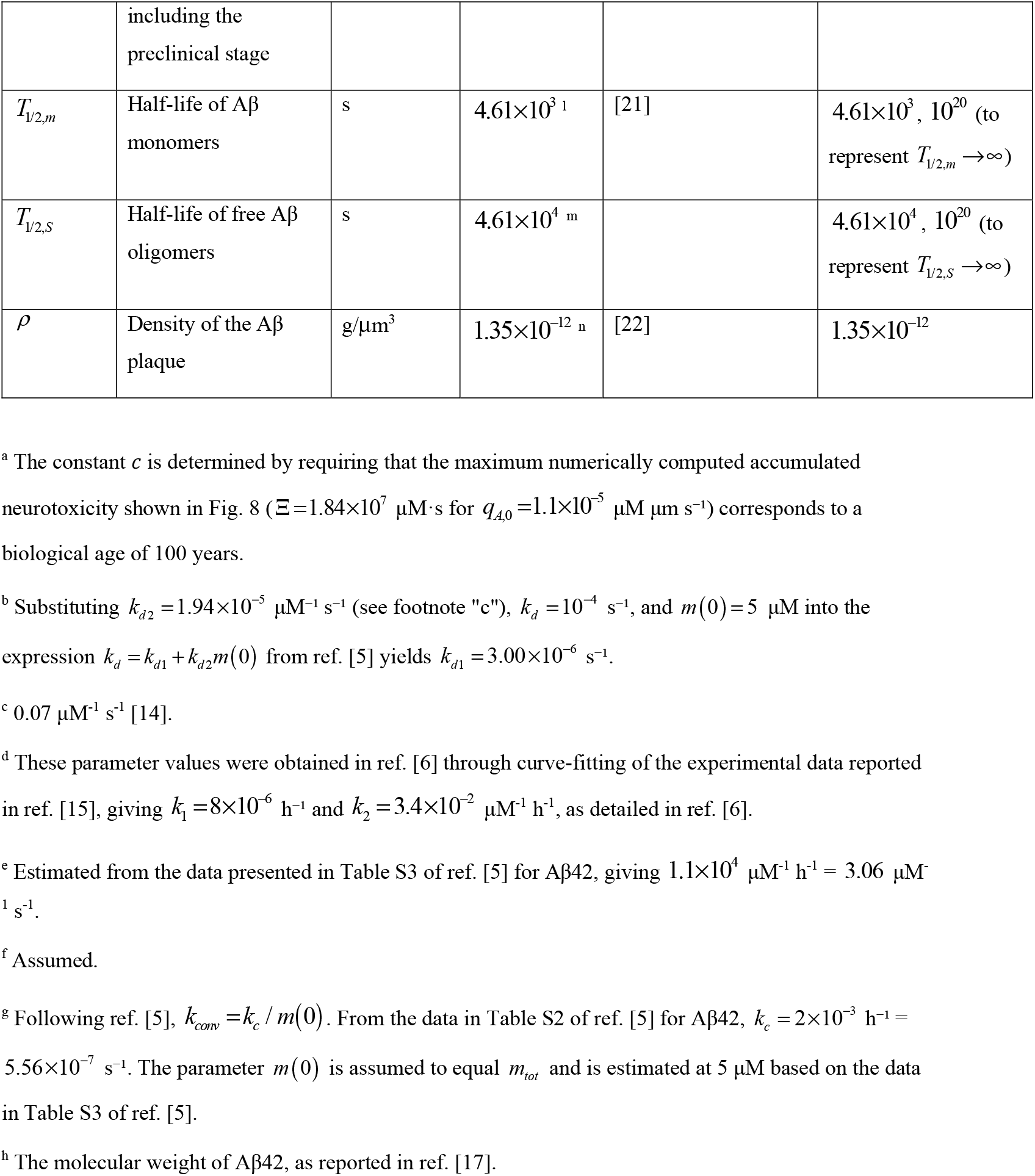
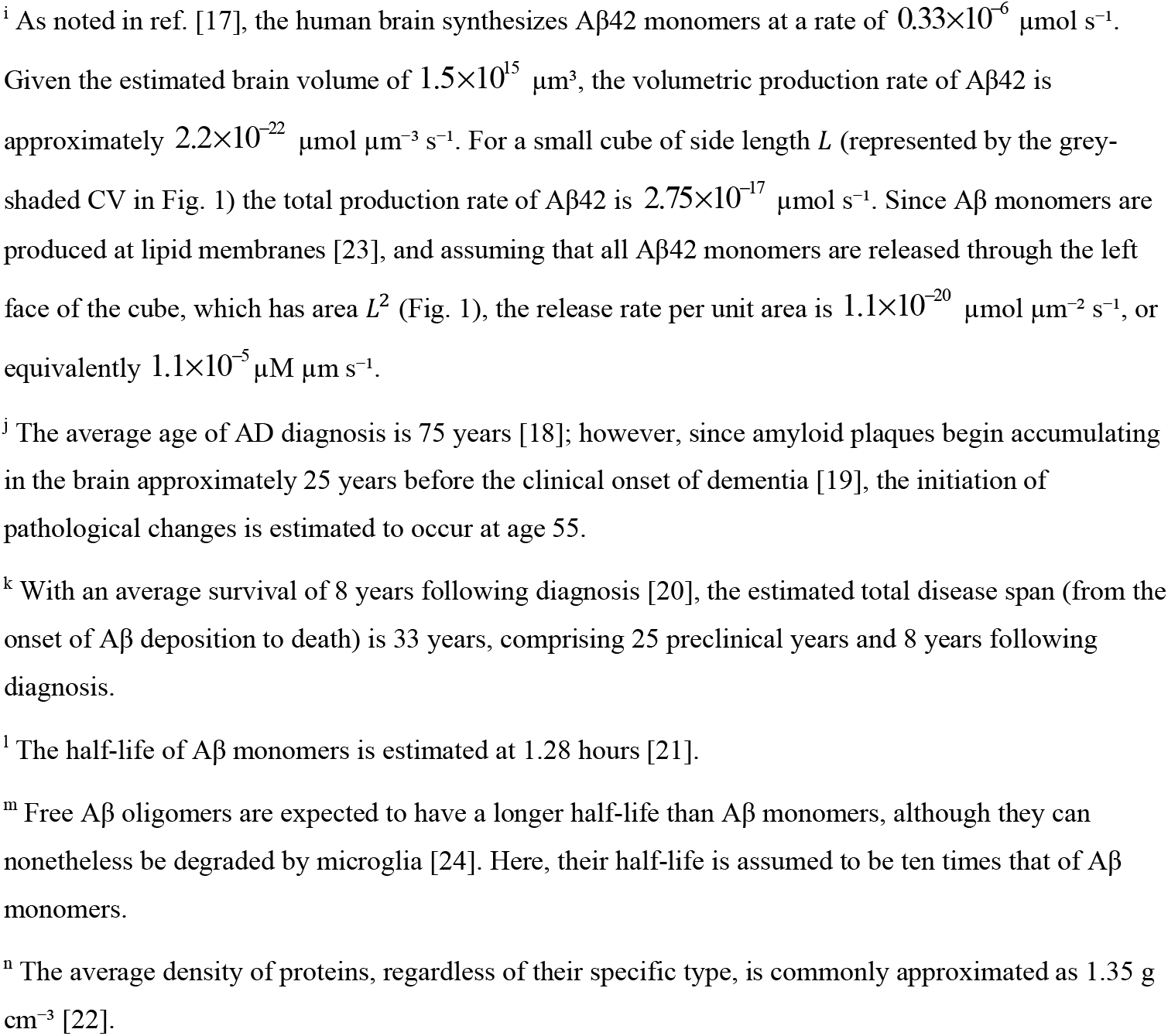
Model parameters.

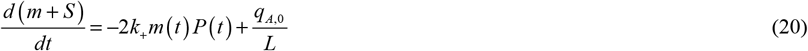

Substituting Eqs. (16)–(18) into Eq. (20) yields:

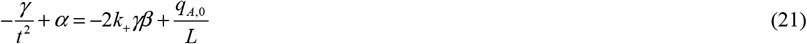

Seeking an approximate solution valid at large times (*t* →∞) and making use of Eq. (19) yields

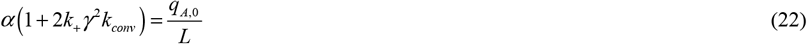

Solving for *α* yields:

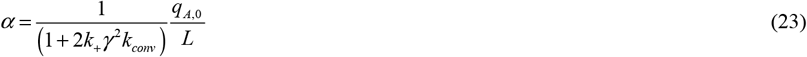

Substituting Eqs. (16)-(18) into Eq. (1) and neglecting the term involving *k*_*conv*_ on account of the small magnitude of *k*_*conv*_ (Table 2) gives:

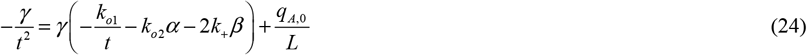

Seeking an approximate solution valid at large times (*t* →∞) yields:

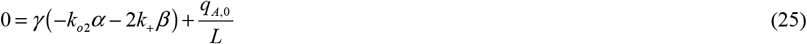

Substituting Eq. (19) into Eq. (25) yields:

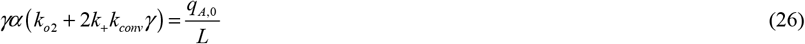

Using Eq. (23) in Eq. (26) yields:

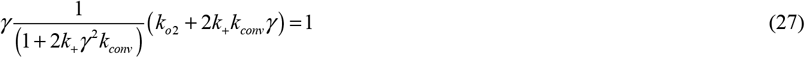

Neglecting the terms involving *k*_*conv*_ on account of the small magnitude of *k*_*conv*_ (Table 2) gives:

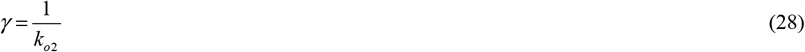

Substituting Eq. (28) into Eq. (23) yields:

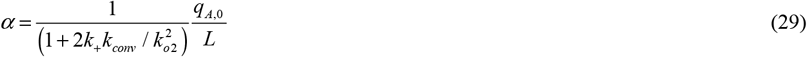

Substituting Eqs. (28) and (29) into Eq. (19), one obtains:

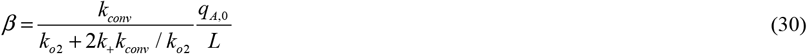

Substituting Eqs. (28)–(30) into Eqs. (16)-(18) gives:

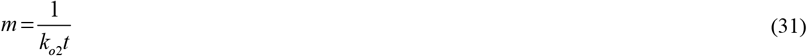

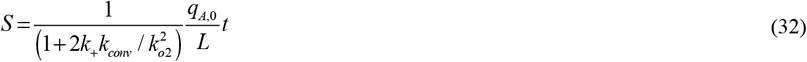

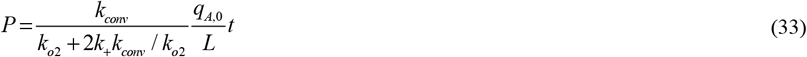

Substituting Eq. (15) into Eq. (10) gives:

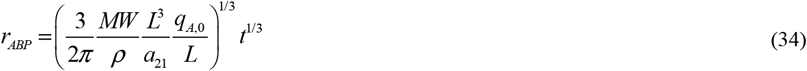

Substituting Eq. (15) into Eq. (11), one obtains:

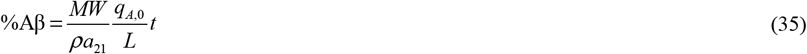

Substituting Eq. (32) into Eq. (12) yields:

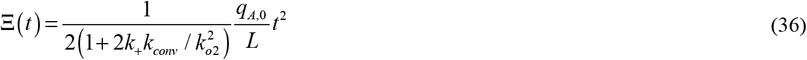

Substituting Eq. (36) into Eq. (13) yields:

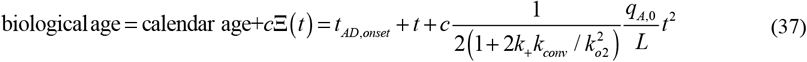

Solving Eq. (35) for *t* and substituting the result into Eq. (36) yields:

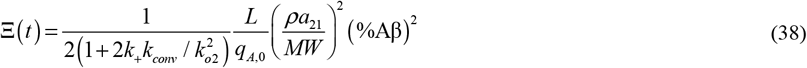

### 2.5. Approximate analytical solution for the limiting case of *k*_−_ = 0, *T*_1/ 2,*m*_ →∞, and *T*_1/ 2,*S*_ →∞ (accounting for oligomer dissociation)

When *k*_*d*1_ and *k*_*d*2_ and are nonzero, adding Eqs. (1), (2), and (4) still yields Eq. (14). Since in this case *m* + *S* ≪ *M* (see Figs. S1b, S2b, and S4b), integrating Eq. (14) again gives Eq. (15). Substituting Eq. (15) into Eqs. (10) and (11) then recovers Eqs. (34) and (35), respectively. Equations (15), (34), and (35) therefore remain valid in the presence of oligomer dissociation.

### 2.6. Approximate analytical solution for the limiting case of *T*_1/ 2,*m*_ →∞ and *T*_1/ 2,*S*_ →∞ (accounting for oligomer dissociation and fibril fragmentation)

When *k*_*d*1_, *k*_*d*2_, and *k*_−_ are nonzero, adding Eqs. (1), (2), and (4) still yields Eq. (14). Since in this case *m* + *S* = *M* (see Figs. S10b, S11b, and S13b), integrating Eq. (14) again gives Eq. (15). It is worth noting that a larger value of *P* (*t*) does not directly affect the accuracy of Eq. (15), since *P* (*t*) does not appear explicitly in Eq. (14). Substituting Eq. (15) into Eqs. (10) and (11) then recovers Eqs. (34) and (35), respectively. Eqs. (15), (34), and (35) therefore remain accurate to a good approximation in the presence of both oligomer dissociation and fibril fragmentation.

## 3. Results

Details of the numerical solution procedure are provided in Section S1 of the Supplementary Materials. Unless stated otherwise in a figure or its caption, all parameter values are those listed in Table 2. The results presented in Figs. 2-10 correspond to the baseline case of negligible oligomer dissociation into monomers (*k*_*d*1_ = 0, *k*_*d*2_ = 0) and negligible fibril fragmentation (*k*_−_ = 0). Throughout, two values are considered for the monomer and oligomer half-lives (*T*_1/ 2,*m*_, and *T*_1/ 2,*S*_): a physiologically relevant value and an infinitely large value representing a complete breakdown of the protein degradation machinery, approximated numerically as 10^20^ s.

The results reveal marked differences between scenarios with intact and impaired degradation machinery. When physiologically relevant half-lives of Aβ monomers and oligomers are used, all species (free monomers, oligomers, fibrils, and plaque) evolve slowly, and their concentrations increase gradually with the monomer production rate *q*_*A*,0_. The free Aβ monomer concentration decreases slowly over time (Fig. 2a), as does the oligomer concentration (Fig. 3a), while the concentration and total mass of fibrillar species increase over time (Figs. 4a and 5a). All these quantities grow with *q*_*A*,0_. When degradation fails, the dynamics change dramatically: the monomer concentration drops rapidly and becomes largely insensitive to *q*_*A*,0_ (Fig. 2b), in agreement with Eq. (31); oligomers accumulate linearly with time (Fig. 3b), in excellent agreement with the analytical solution of Eq. (32), and fibrillar species grow at a rate approximately three orders of magnitude higher than in the intact degradation scenario, consistent with the analytical predictions of Eqs. (33) and (15) (Figs. 4b and 5b).

**Fig. 3.**
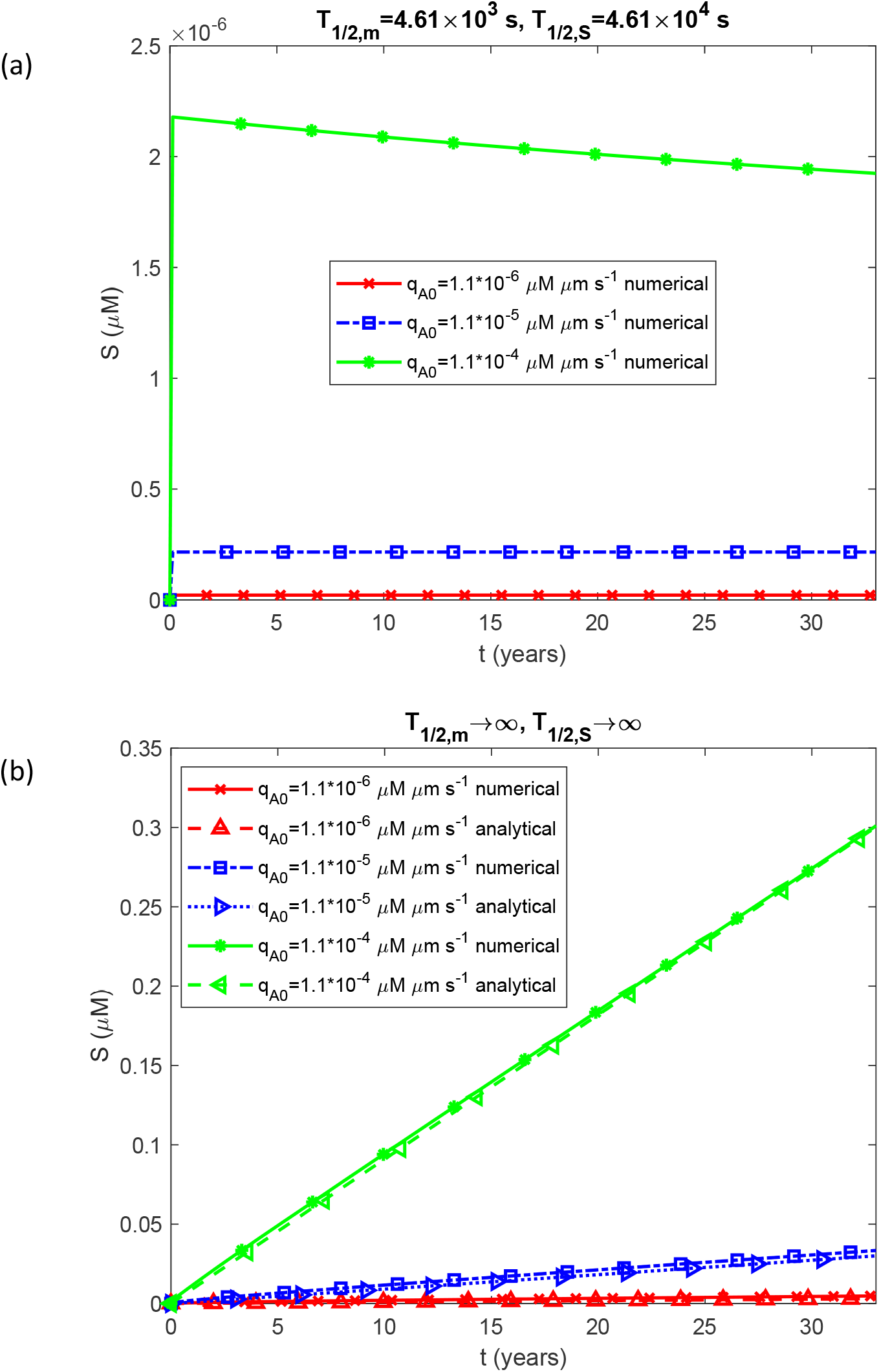
Molar concentration of free on-pathway Aβ oligomers as a function of time, *S* (*t*). The case of negligible oligomer dissociation into monomers (*k*_*d*1_ = 0, *k*_*d*2_ = 0) and negligible fibril fragmentation (*k*_−_ = 0).

**Fig. 4.**
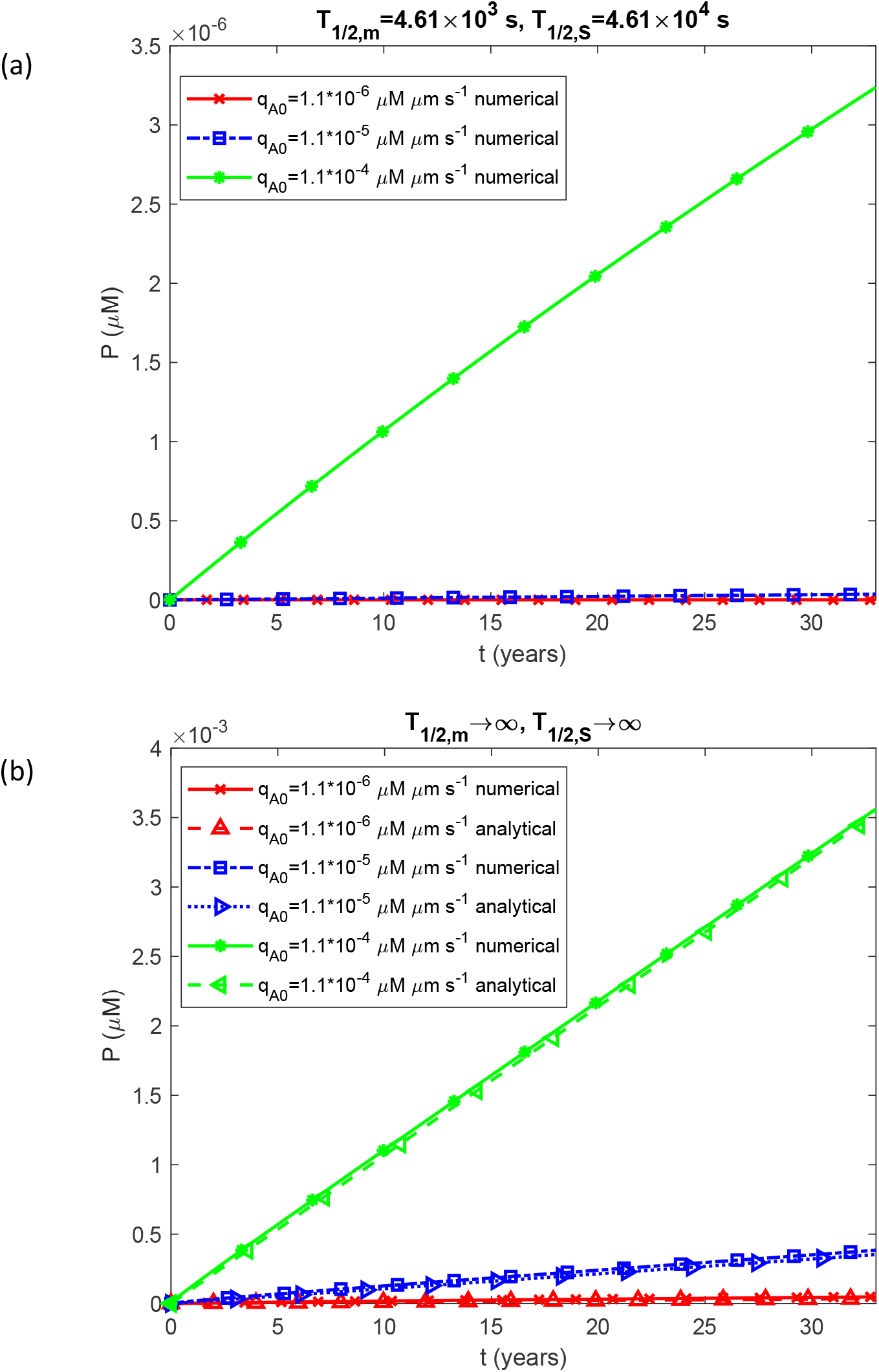
Molar concentration of Aβ fibrillar species of varying length as a function of time, *P*(*t*). The case of negligible oligomer dissociation into monomers (*k*_*d*1_ = 0, *k*_*d*2_ = 0) and negligible fibril fragmentation (*k*_−_ = 0).

**Fig. 5.**
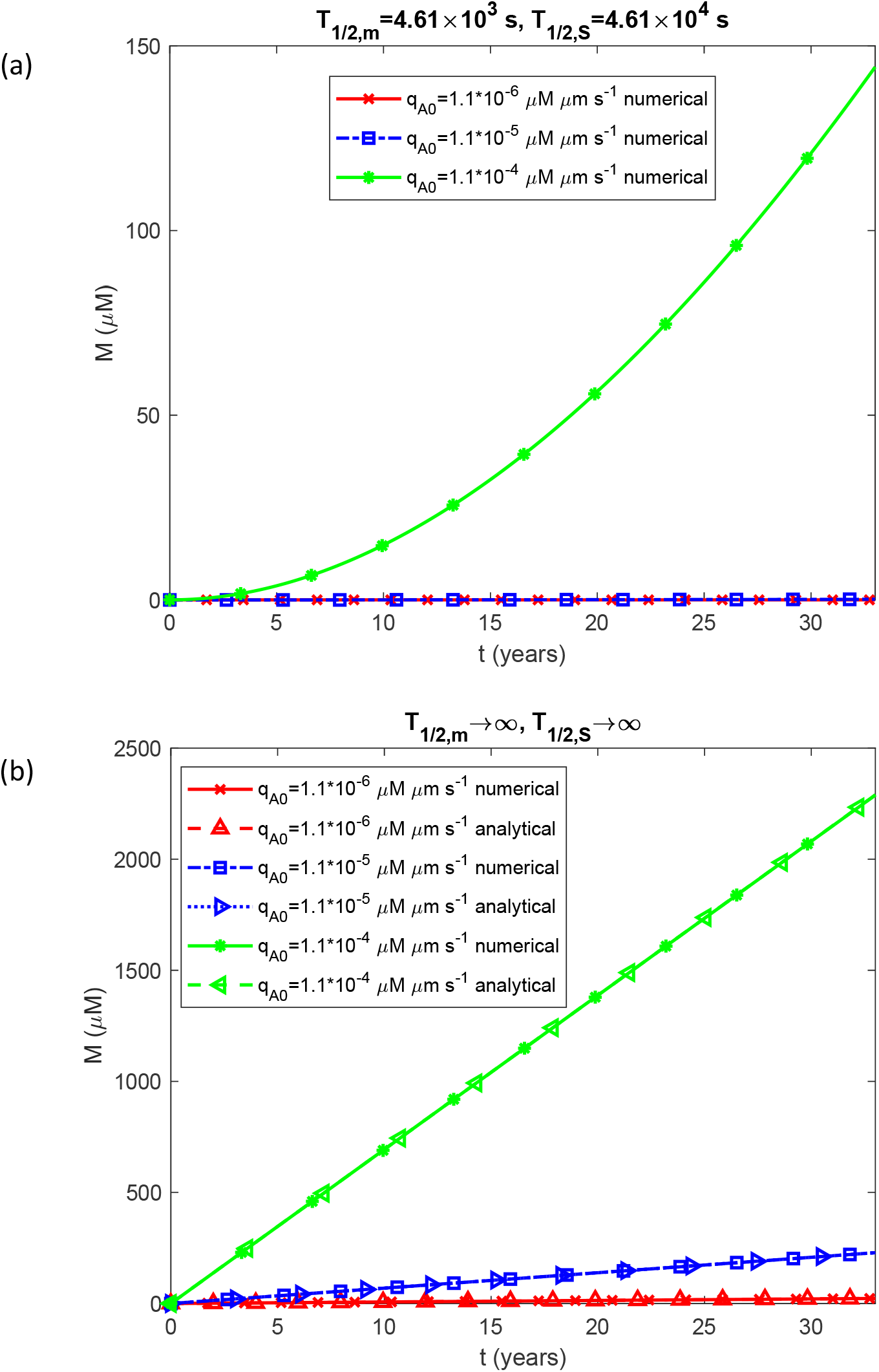
Molar concentration of Aβ monomers incorporated into fibrillar species of varying length as a function of time, *M* (*t*). The case of negligible oligomer dissociation into monomers (*k*_*d*1_ = 0, *k*_*d*2_ = 0) and negligible fibril fragmentation (*k*_−_ = 0).

These molecular-level changes manifest directly in plaque growth. For physiologically relevant half-lives, both the plaque radius and the fraction of the CV occupied by the plaque increase gradually with time and with *q*_*A*,0_ (Figs. 6a and 7a). Under impaired degradation, the plaque radius scales with the cube root of time, as predicted by Eq. (34), while the occupied volume fraction grows linearly with time, consistent with Eq. (35). This linear growth is a direct consequence of the constant monomer production rate assumed in the model: in the absence of degradation, monomers accumulate unimpeded and are progressively converted into plaque.

**Fig. 6.**
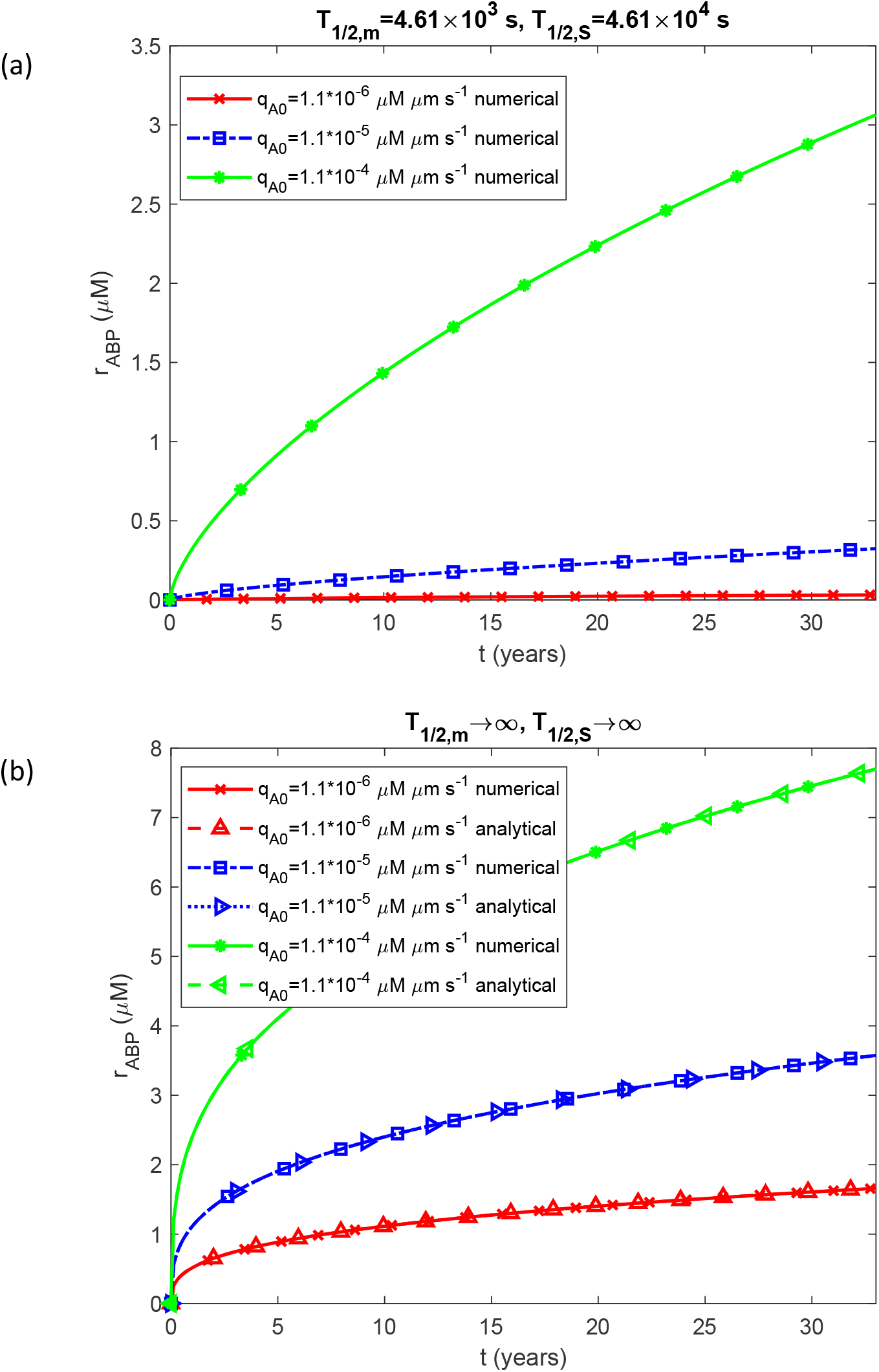
Radius of the Aβ plaque as a function of time, *r*_*ABP*_ (*t*). The case of negligible oligomer dissociation into monomers (*k*_*d*1_ = 0, *k*_*d*2_ = 0) and negligible fibril fragmentation (*k*_−_ = 0).

**Fig. 7.**
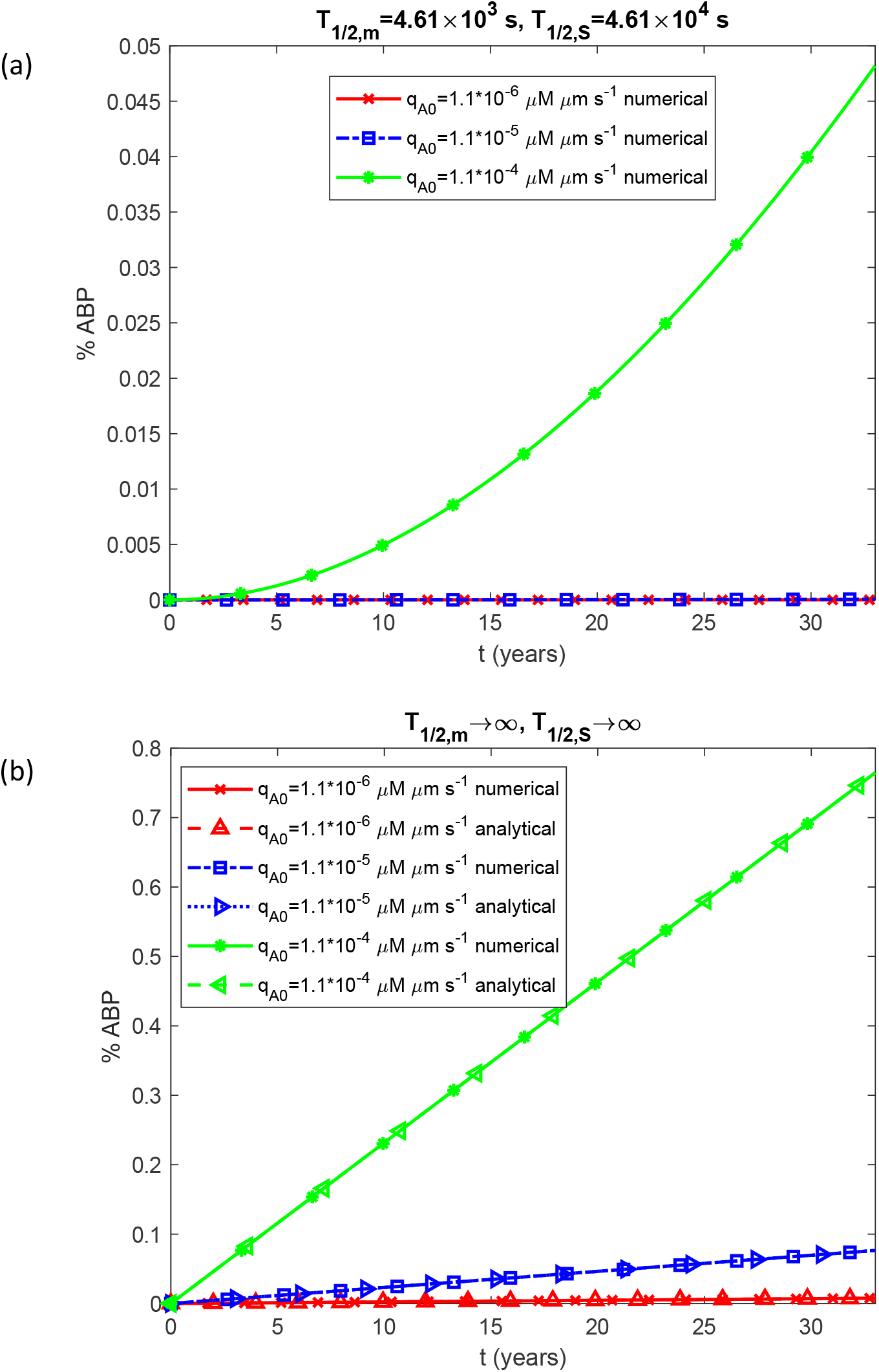
Percentage of the CV occupied by the Aβ plaque as a function of time. The case of negligible oligomer dissociation into monomers (*k*_*d*1_ = 0, *k*_*d*2_ = 0) and negligible fibril fragmentation (*k*_−_ = 0).

Under physiological conditions, accumulated neurotoxicity grows linearly with time, and biological age closely tracks calendar age (Figs. 8a and 9a). When degradation is impaired, accumulated neurotoxicity grows as the square of time, in agreement with Eq. (36), and biological age advances considerably faster than calendar age, particularly at high monomer production rates (Fig. 9b).

**Fig. 8.**
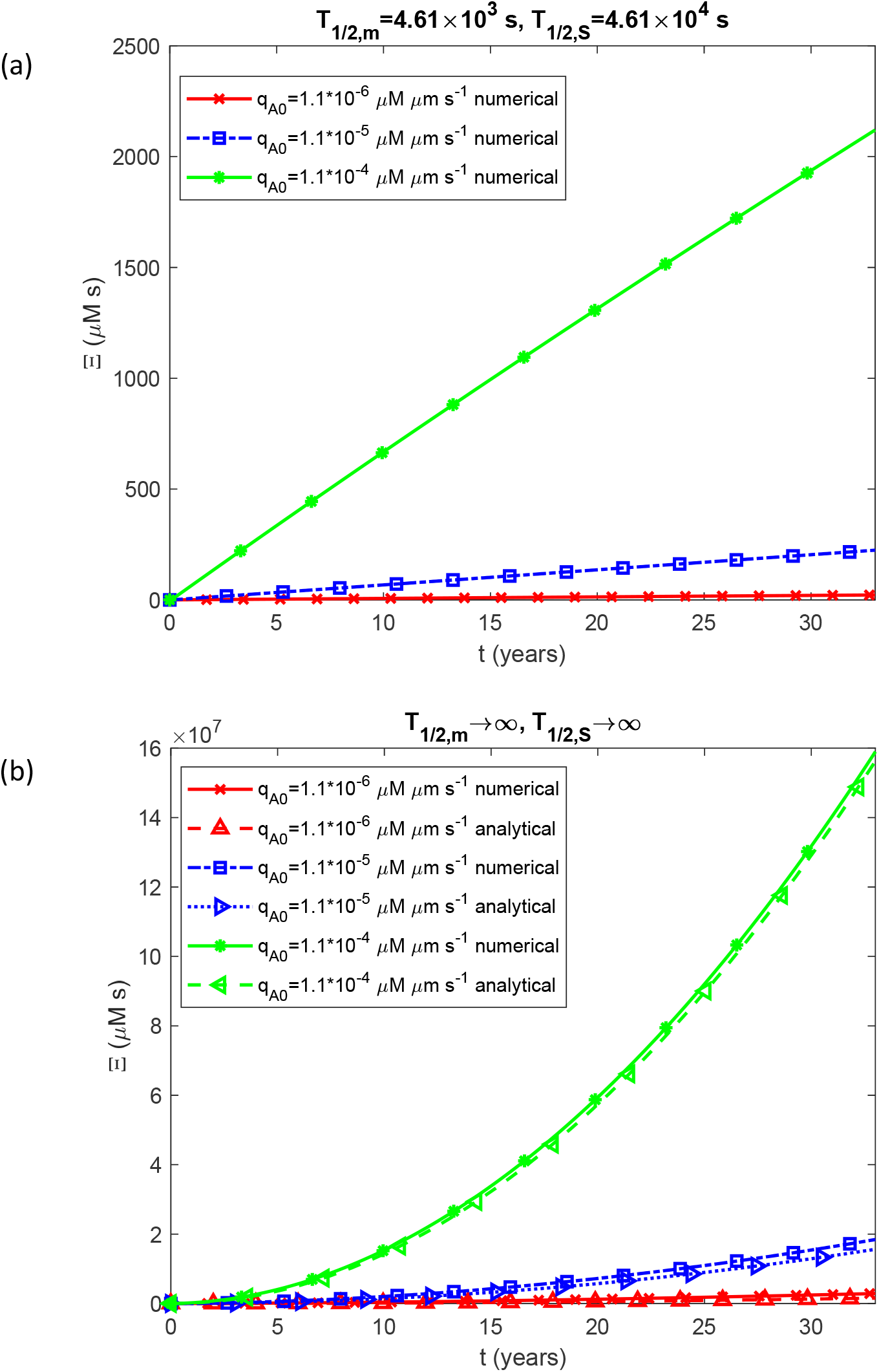
Accumulated neurotoxicity as a function of time, Ξ(*t*). The case of negligible oligomer dissociation into monomers (*k*_*d*1_ = 0, *k*_*d*2_ = 0) and negligible fibril fragmentation (*k*_−_ = 0).

**Fig. 9.**
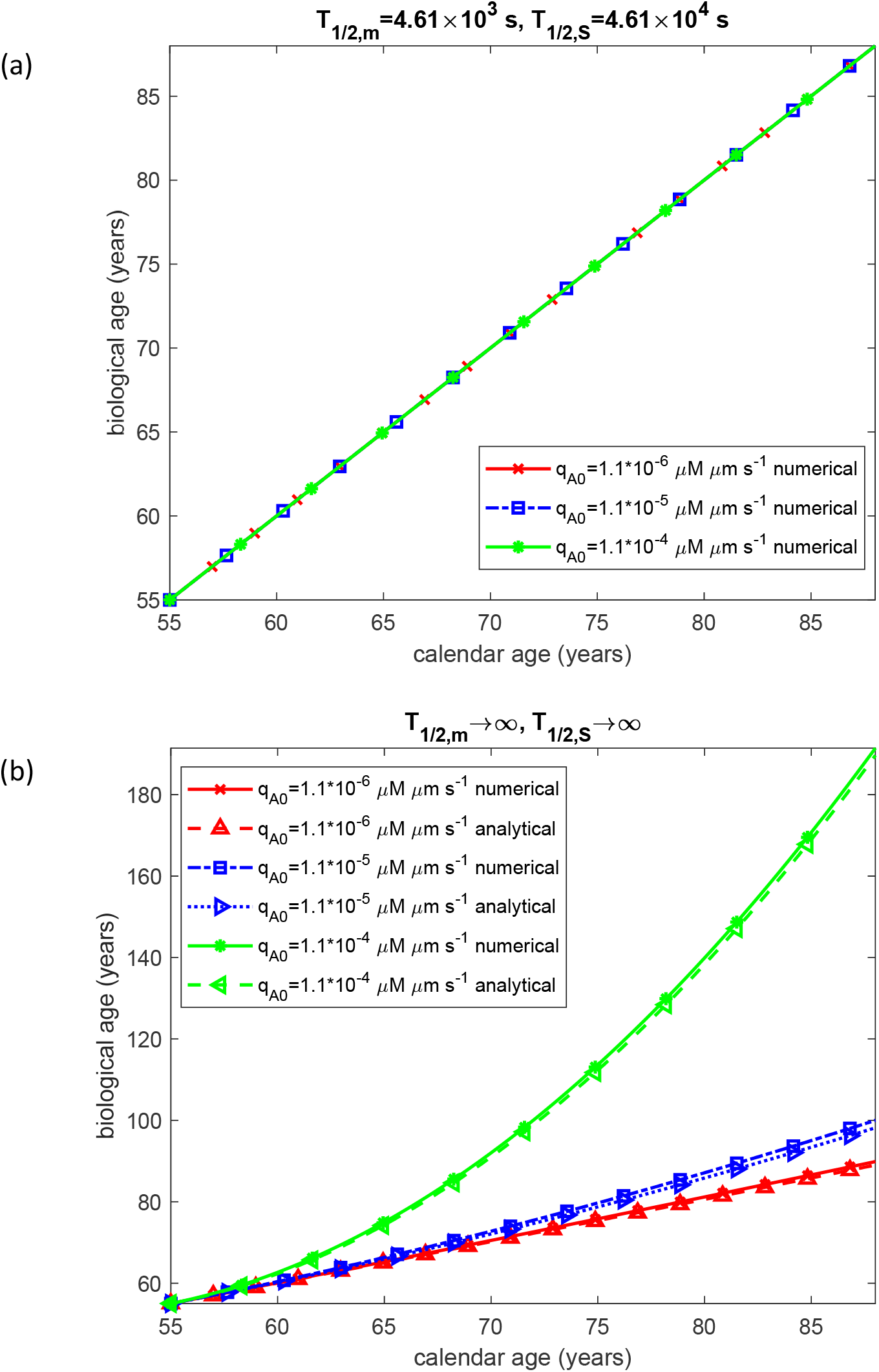
Biological age as a function of calendar age. The case of negligible oligomer dissociation into monomers (*k*_*d*1_ = 0, *k*_*d*2_ = 0) and negligible fibril fragmentation (*k*_−_ = 0).

It is worth noting that for the case of impaired Aβ degradation machinery, the analytical solution for oligomer concentration, Eq. (32), predicts a linear dependence of *S* (*t*) on time, which is confirmed by the numerical solutions shown in Fig. 3b. This linear growth in turn gives rise to the quadratic dependence of accumulated neurotoxicity on time given by Eq. (36) and illustrated in Fig. 8b. A practical implication of this result is that calculating accumulated neurotoxicity requires only a single measurement of oligomer concentration combined with knowledge of the individual’s calendar age, since the integral 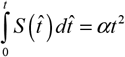 (see Eq. (12)), and the constant *α* can be determined from that single measurement. Techniques for quantifying Aβ oligomer concentrations in human brain tissue have been reported [35], though these currently require post-mortem or biopsy samples.

For *q*_*A*,0_ = 1.1×10^−4^ μM μm s^−1^, the model predicts that biological age reaches 100 years at a calendar age of only approximately 72 years. The relationship between accumulated neurotoxicity and plaque volume further reveals a power-law dependence: on a log-log plot this relationship appears as a straight line with slope 2, as predicted analytically by Eq. (38), indicating that neurotoxicity scales with the square of the plaque volume fraction (Fig. 10b).

**Fig. 10.**
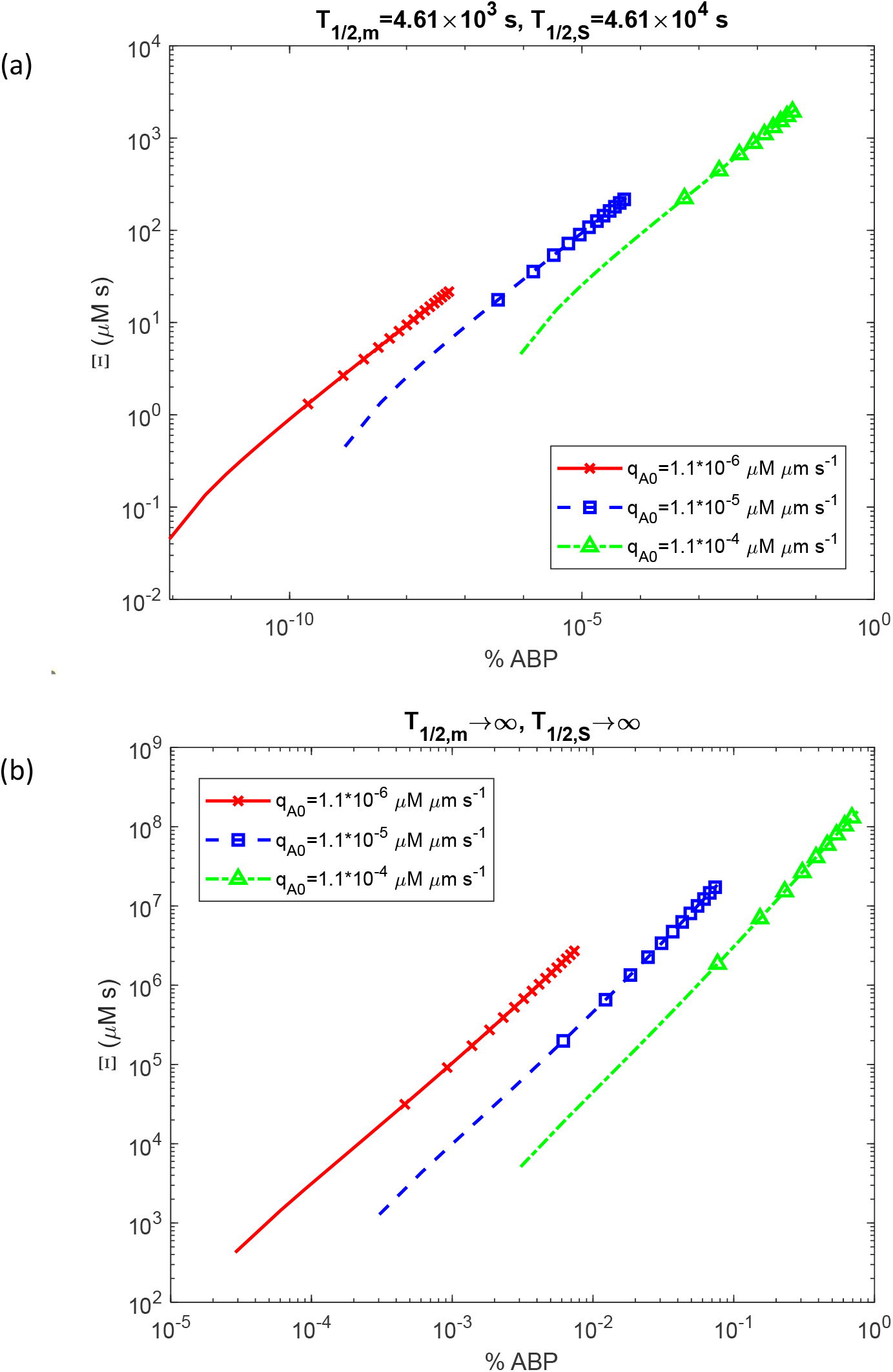
Accumulated neurotoxicity as a function of the percentage of the CV occupied by the Aβ plaque. The case of negligible oligomer dissociation into monomers (*k*_*d*1_ = 0, *k*_*d*2_ = 0) and negligible fibril fragmentation (*k*_−_ = 0).

The results presented in Figs. S1-S9 correspond to the case in which oligomer dissociation into monomers is accounted for (*k*_*d*1_ = 3×10^−6^ s^-1^, *k*_*d*2_ = 1.94 × 10^−5^ μM^-1^ s^-1^) while fibril fragmentation is neglected (*k*_−_ = 0). The key distinction introduced by this variant of the model is that monomers can be regenerated through oligomer dissociation, a process that can itself be catalyzed by fibril surfaces. As shown below, this additional pathway has profound consequences for the dynamics of every species in the system.

The most immediate effect of oligomer dissociation is visible at the level of the free monomer and oligomer pools. Rather than decreasing monotonically as in the baseline case, the free Aβ monomer concentration now rapidly increases to a steady-state value and thereafter remains constant, scaling with the monomer production rate under both physiological (Fig. S1a) and impaired (Fig. S1b) degradation conditions. The steady-state monomer concentration under impaired degradation is notably higher than in the corresponding baseline case (cf. Fig. 2b), reflecting the additional source of monomers provided by oligomer dissociation. The oligomer pool, by contrast, is depleted more rapidly than in the baseline scenario: its concentration decreases over time under physiological conditions (Fig. S2a) and drops sharply when degradation is impaired (Fig. S2b), in contrast to the accumulation observed in Fig. 3b. Interestingly, under impaired degradation, the oligomer concentration decreases with increasing monomer production rate (Fig. S2b), a counterintuitive behavior driven by the enhanced fibril-catalyzed dissociation that accompanies higher monomer flux.

These upstream changes propagate into fibril formation and plaque growth. The concentration of fibrillar species of varying length increases with time and with the monomer production rate (Fig. S3a), but an order of magnitude more slowly than in the absence of oligomer dissociation (cf. Fig. 4a). Under impaired degradation, the fibrillar concentration reaches a steady-state value which is approximately three orders of magnitude lower than the largest value observed in the baseline case (cf. Fig. 4b), because oligomer dissociation continuously diverts oligomeric species away from the fibril formation pathway. Consistently, the total fibril mass grows approximately five times more slowly than in the baseline scenario under physiological conditions (cf. Figs. S4a and 5a), while under impaired degradation it increases linearly with time in exact agreement with the path predicted by Eq. (15) (Fig. S4b). The plaque radius and the volume fraction of the CV occupied by the plaque exhibit the same trend: both grow with time and with the monomer production rate, but roughly 1.5 and 5 times more slowly, respectively, than in the baseline case under physiological conditions (cf. Figs. S5a, S6a with Figs. 6a, 7a), and both follow the analytical solutions of Eqs. (34) and (35) precisely when degradation is impaired (Figs. S5b and S6b).

The inclusion of oligomer dissociation also has significant implications for neurotoxicity and biological aging. Under physiological conditions, accumulated neurotoxicity increases over time at a rate approximately one order of magnitude lower than in the baseline case (cf. Figs. S7a and 8a), and grows with the monomer production rate. Under impaired degradation, accumulated neurotoxicity continues to rise but at a rate approximately six orders of magnitude lower than in the absence of oligomer dissociation (cf. Figs. S7b and 8b); moreover, it decreases as the monomer production rate increases, a further consequence of the dissociation dynamics described above. Because accumulated neurotoxicity remains low throughout the simulated lifespan, biological age and calendar age remain nearly indistinguishable up to a calendar age of 88 years in both degradation scenarios (Fig. S8a,b). Finally, the relationship between accumulated neurotoxicity and plaque burden no longer follows a simple power law: unlike the baseline case, the log-log plot is no longer linear (Fig. S9a,b), and accumulated neurotoxicity reaches values several orders of magnitude lower than in the absence of oligomer dissociation, most dramatically so under impaired degradation (cf. Figs. S9b and 10b). Taken together, these results demonstrate that oligomer dissociation could act as a powerful protective mechanism, substantially slowing fibril growth, plaque expansion, and the accumulation of neurotoxic burden.

The results presented in Figs. S10-S18 correspond to the case in which both oligomer dissociation into monomers and fibril fragmentation are accounted for (*k*_*d*1_ = 3×10^−6^ s^-1^, *k*_*d*2_ = 1.94 ×10^−5^ μM^-1^ s^-1^, *k*_−_ = 10^−14^ s^-1^). The distinguishing feature of this scenario is that new fibrils can be generated through the fragmentation of existing ones, introducing an additional mechanism that alters the dynamics of every species in the system.

The most immediate consequence of fibril fragmentation is seen at the level of monomer consumption. Unlike the case with oligomer dissociation alone, where the free Aβ monomer concentration rapidly reaches a steady state, fragmentation continuously generates new fibril ends that serve as elongation sites, thereby increasing the rate at which monomers are consumed. As a result, the monomer concentration decreases monotonically over time under both physiological (Fig. S10a) and impaired (Fig. S10b) degradation conditions, rather than plateauing as observed in the absence of fragmentation (cf. Figs. S1a and S1b). The oligomer concentration decreases more rapidly than in the case without fibril fragmentation (cf. Figs. S11a,b and S2a,b), because fragmentation diverts monomers into fibril elongation, accelerating the depletion of the soluble oligomer pool.

The effects of fragmentation on fibril concentration are substantial. Under physiological conditions, the concentration of fibrillar species of varying length increases with time and with the monomer production rate (Fig. S12a), reaching values approximately four orders of magnitude greater than in the fragmentation-free case (cf. Fig. S3a), and the total fibril mass grows approximately 50-fold faster (cf. Figs. S13a and S4a). Under impaired degradation, the fibrillar concentration increases continuously with time and with the monomer production rate (Fig. S12b), in marked contrast to the fragmentation-free scenario where it approached a steady state (cf. Fig. S3b), while the total fibril mass increases linearly with time in agreement with the analytical solution of Eq. (15) (Fig. S13b).

These upstream changes translate into markedly accelerated plaque growth under physiological conditions: the plaque radius grows approximately four-fold faster than in the fragmentation-free case (cf. Figs. S14a and S5a), and the fraction of the CV occupied by the plaque increases approximately 50-fold faster (cf. Figs. S15a and S6a), with both quantities scaling with the monomer production rate. Under impaired degradation, the plaque radius grows in proportion to the cube root of time, in excellent agreement with Eq. (34) (Fig. S14b), and the occupied volume fraction increases linearly with time, consistent with Eq. (35) (Fig. S15b).

Despite the accelerated plaque growth observed under physiological conditions, the effect of fibril fragmentation on neurotoxicity is, counterintuitively, attenuating rather than amplifying. Accumulated neurotoxicity increases with time and with the monomer production rate under both physiological and impaired degradation conditions (Figs. S16a,b), but does so approximately two-fold more slowly than in the case without fragmentation (cf. Figs. S7a,b). This attenuation arises because fragmentation diverts monomers into fibril elongation, thereby reducing the pool of soluble oligomers that drive neurotoxic burden. Consequently, accumulated neurotoxicity is even lower than in the case of oligomer dissociation alone, and biological age remains practically identical to calendar age throughout the entire simulated lifespan (Figs. S17a,b). The relationship between accumulated neurotoxicity and plaque burden likewise shows no simple power-law dependence: the log-log plot remains nonlinear (Figs. S18a,b), reflecting the complex and indirect coupling between plaque volume and neurotoxic burden in this scenario.

## 4. Discussion, limitations, and future directions

The modelling results are consistent with experimental and clinical observations reported in the literature, and help to shed light on some of these observations while also placing them in a therapeutic context.

Central to the model is the assumption that soluble Aβ oligomers are the primary neurotoxic species. The model defines accumulated neurotoxicity in terms of the time-integrated concentration of soluble Aβ oligomers, rather than plaque burden. This choice is supported by a substantial body of experimental literature. As early as 1999, ref. [27] demonstrated that soluble Aβ concentration is elevated threefold in AD and correlates far more closely with markers of disease severity than does plaque count. Subsequent work has identified multiple mechanisms by which oligomers exert neurotoxic effects, including direct interactions with synaptic receptors that contribute to progressive synaptic failure [28-31]. More specifically, ref. [36] proposed that soluble Aβ oligomers partially reduce NMDAR-mediated calcium influx in the hippocampus, favoring the induction of long-term depression pathways and thereby contributing to synapse loss. Ref. [37] further showed that prolonged exposure of cortical neurons to Aβ oligomers causes NADPH oxidase-mediated production of reactive oxygen species and mitochondrial dysfunction, impairing NMDA receptor function and promoting neuronal cytotoxicity. Ref. [38] noted that accumulation of soluble Aβ oligomers, which begins one to two decades before the appearance of clinical symptoms, induces neuronal hyperactivity that in turn triggers synapse dysfunction and loss. The present model is consistent with this picture: accumulated neurotoxicity builds up gradually over decades, with the rate of accumulation depending critically on the integrity of the protein degradation machinery rather than on plaque volume alone.

The model further demonstrates that neural damage is not determined solely by conditions at any particular moment in time, but is instead path-dependent. A key conceptual feature of the model is that biological age is governed by the time-integrated oligomer burden rather than by the instantaneous plaque burden, reflecting a longitudinal rather than cross-sectional dependence. This is consistent with the clinical observation, reported by ref. [39], that AD patients with similar single-timepoint biomarker values can exhibit markedly divergent cognitive trajectories, supporting the view that neural damage in AD is path-dependent and determined by the historical trajectory of biomarker accumulation rather than solely by current plaque burden. The model formalizes this notion by defining accumulated neurotoxicity, which encodes the entire history of oligomer exposure. This perspective is further supported by ref. [33], who found that the yearly rate of change in Aβ burden improves the prediction of cognitive decline beyond what a single static measurement can provide.

A substantial body of literature has established a relationship between Aβ plaque burden and neural damage and cognitive decline in AD [40-43]. If accumulated neurotoxicity, defined as the time integral of the Aβ oligomer concentration, is the true determinant of neural damage in AD, then a relationship between accumulated neurotoxicity and Aβ plaque burden must exist.

In the limiting case of impaired degradation machinery, the model predicts that this relationship takes the form of a power law, with accumulated neurotoxicity scaling as the square of the plaque volume fraction. However, this relationship breaks down when oligomer dissociation and fibril fragmentation are included, and the log-log plot becomes nonlinear. This is consistent with the clinical evidence that plaque burden alone is an imperfect surrogate for disease severity. Ref. [3] found that the ratio of Aβ oligomer levels to plaque density fully distinguished demented from non-demented patients, with no overlap between groups in this derived variable, demonstrating that two patients with equivalent plaque burdens can have very different cognitive outcomes depending on their respective levels of soluble Aβ oligomers. This is precisely what the model would predict: because neurotoxicity is driven by integrated oligomer exposure, two individuals with identical plaque burdens but different oligomer dynamics (determined, for instance, by differences in degradation efficiency) will accumulate very different neurotoxic burdens over time.

The modelling results carry significant implications for the development of AD therapies. In particular, the model’s prediction that plaque clearance alone cannot reverse the neurotoxic burden already accumulated over time is consistent with the clinical trial record. A drug withdrawal trial in AD subjects who had completed 78 weeks of lecanemab treatment reported progressive cognitive decline despite a persistently low plaque burden, indicating that when active treatment is not maintained, the toxicity of soluble Aβ oligomers remains unabated even in the near-absence of Aβ plaques [2]. More broadly, ref. [2] noted that only agents targeting soluble Aβ oligomers have shown clinical efficacy, while agents that predominantly target monomers or plaques have failed to demonstrate clinical benefit. These observations reinforce the model’s emphasis on the oligomer pool as the therapeutically relevant quantity, and suggest that effective intervention must reduce oligomer exposure throughout the disease course rather than simply reducing plaque burden at a single point in time. The model further predicts that impairment of the protein degradation machinery dramatically accelerates oligomer accumulation and neurotoxic burden, advancing biological age well ahead of calendar age. This underscores the potential value of therapeutic strategies aimed at restoring or enhancing proteolytic clearance of Aβ species.

The theoretically predicted plaque growth kinetics are supported by published experimental data. Aβ burden assessed by PET imaging has been shown to increase approximately linearly at a rate of 0.043 SUVR per year over roughly 19 years before slowing and approaching a plateau [44]. The present model predicts linear growth of plaque volume with time in the limiting case of impaired degradation, and a slower, sub-linear growth under physiologically relevant degradation conditions. Although some studies report a sigmoidal pattern of overall plaque burden growth [45], [1], the prolonged linear phase of approximately 20 years reported in humans is broadly consistent with the model’s predictions during the period of active plaque growth, before saturation effects become dominant.

Several simplifying assumptions limit the present model and point to directions for future work. The accumulated neurotoxicity criterion proposed here operates at the local cellular level. Extending it to encompass different brain regions would require a model capable of describing the propagation of oligomeric and fibrillar Aβ species throughout the entire brain. The model also treats the Aβ monomer production rate as constant, whereas in reality it likely varies with neuronal activity, membrane composition, and disease stage. Furthermore, the present model does not account for the role of tau pathology, neuroinflammation, or synaptic loss, all of which interact with Aβ dynamics in ways that may significantly modulate neurotoxic outcomes. Finally, the accumulated neurotoxicity framework, while conceptually motivated, is a simplification of the complex and incompletely understood cascade of events linking oligomer exposure to cognitive decline. Incorporating more detailed mechanistic descriptions of synaptic dysfunction and neuronal loss into the model remains an important goal for future work. The lumped capacitance approximation, which treats all Aβ concentrations as spatially uniform within the CV, neglects the spatial heterogeneity of plaque distribution in neural tissue and the diffusion of Aβ species between plaques. Future models could incorporate spatial gradients and inter-plaque transport to capture the propagation of Aβ pathology across brain regions.

## Abbreviations

AD: Alzheimer’s disease
Aβ: amyloid beta
CV: control volume

## Author Contribution Statement

AVK is the sole author of this paper.

## Funding Data

National Science Foundation (Grant No. DMS-2451660; Funder ID: 10.13039/100000146).

Alexander von Humboldt Foundation through the Humboldt Research Award (Funder ID: 10.13039/100005156).

## Data Availability Statement

This article has no additional data.

## Ethics Statement

None.

## Supplemental Materials

### S1. Numerical Solution

The system of differential equations, Eqs. (1)-(4), subject to the initial conditions of Eq. (5), was integrated numerically using the ODE45 solver in MATLAB (R2024a, MathWorks, Natick, MA, USA). This solver employs an adaptive step-size Runge–Kutta scheme and was selected for its reliability and accuracy. To guarantee high numerical precision, both the relative and absolute tolerances (RelTol and AbsTol) were set to 10^−10^.

### S2. Supplementary figures

**Fig. S1.**
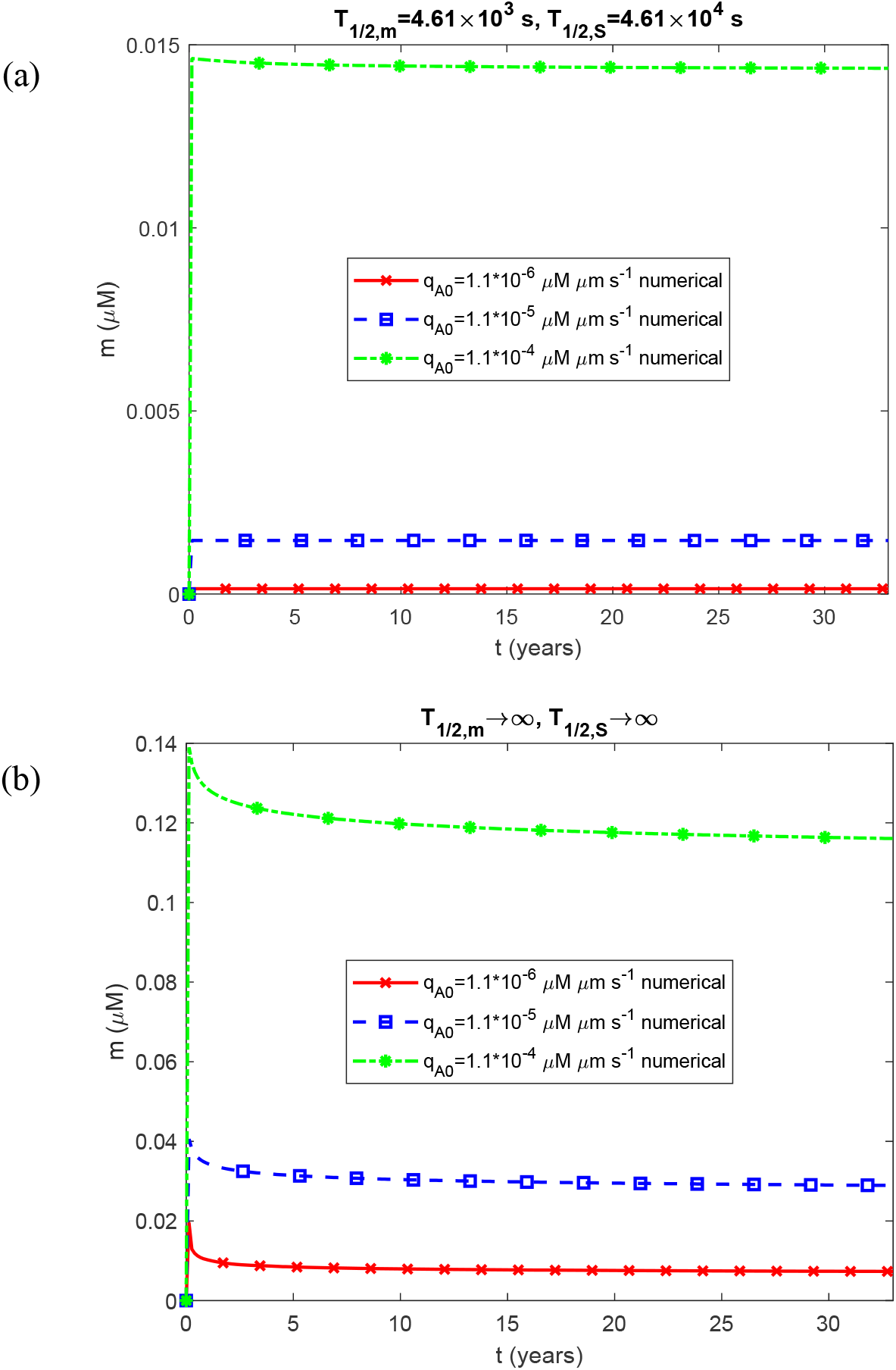
Molar concentration of free Aβ monomers as a function of time, *m*(*t*). The case in which oligomer dissociation into monomers is accounted for (*k*_*d*1_ = 3×10^−6^ s^-1^, *k*_*d*2_ = 1.94 ×10^−5^ μM^-1^ s^-1^) and fibril fragmentation is neglected (*k*_−_ = 0).

**Fig. S2.**
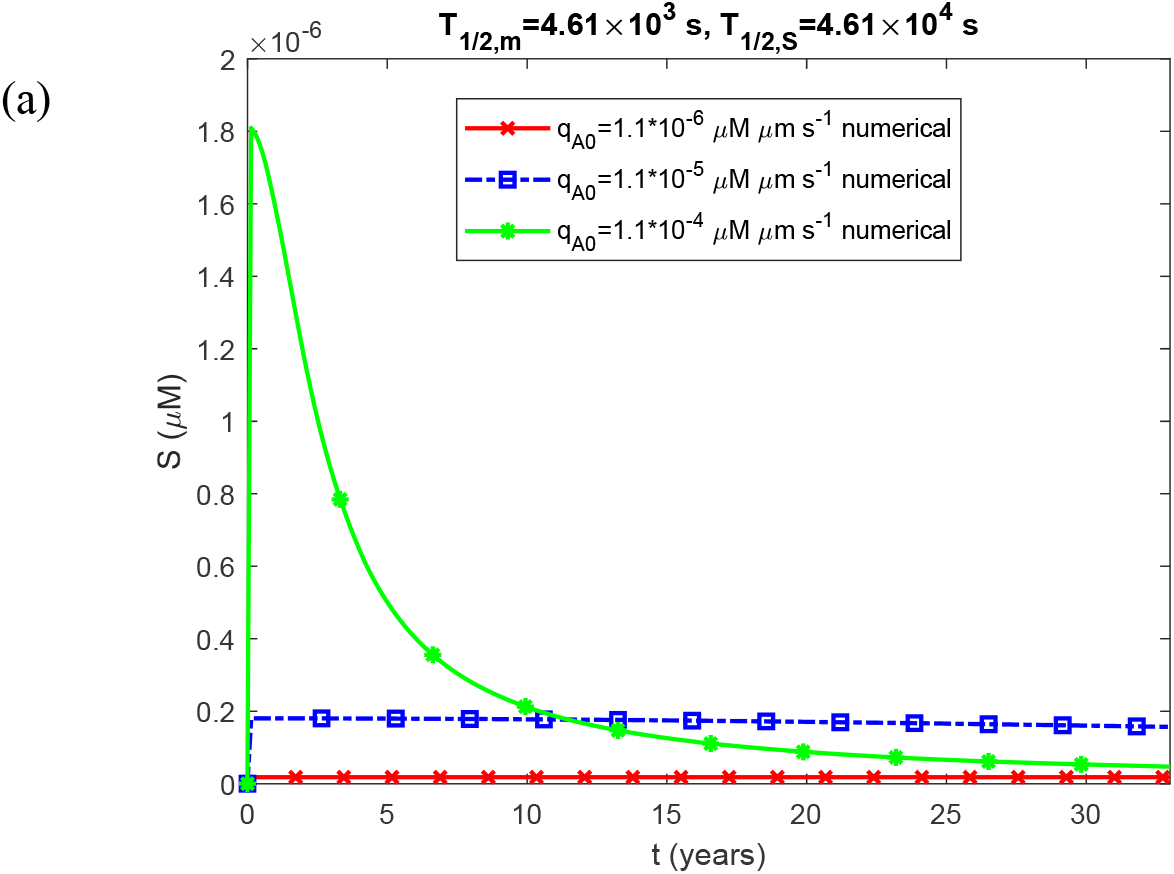
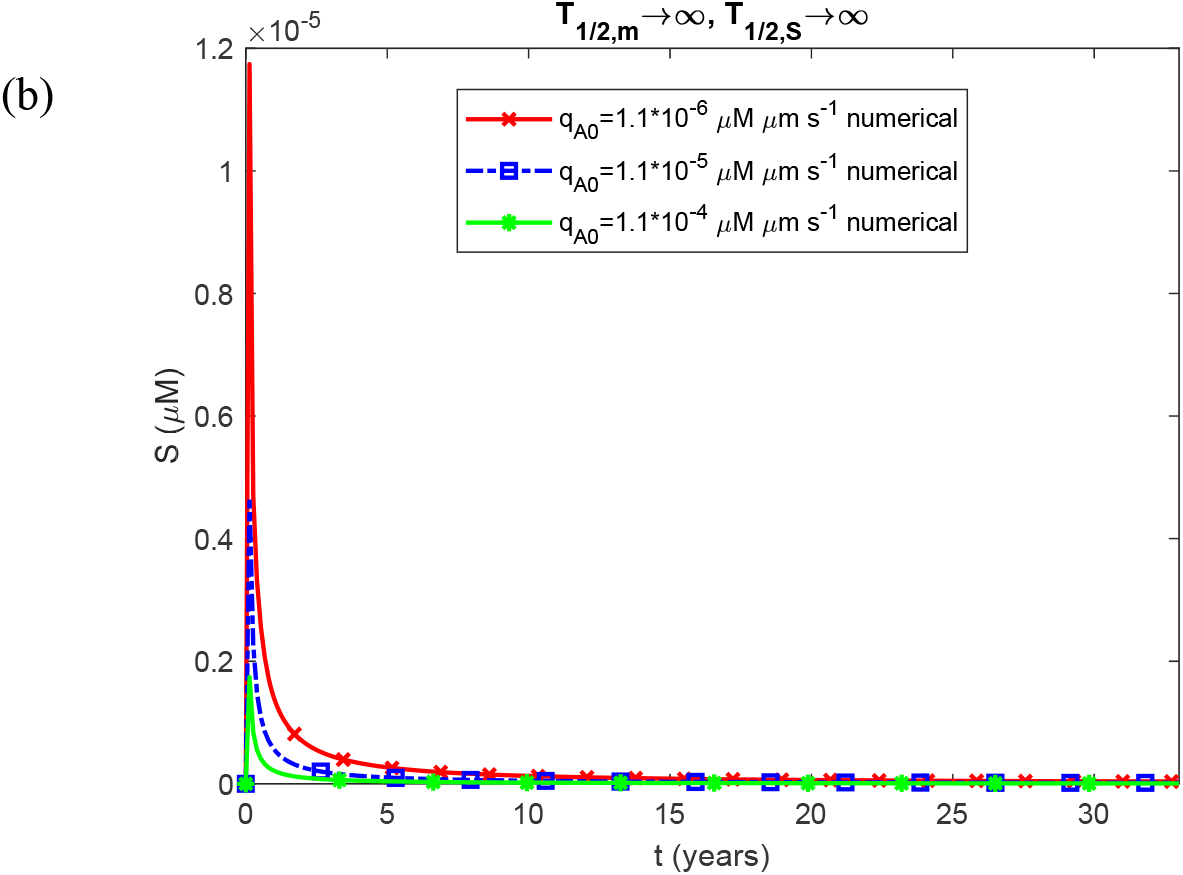
Molar concentration of free on-pathway Aβ oligomers as a function of time, *S* (*t*). The case in which oligomer dissociation into monomers is accounted for (*k*_*d*1_ = 3×10^−6^ s^-1^, *k*_*d*2_ = 1.94 ×10^−5^ μM^-1^ s^-1^) and fibril fragmentation is neglected (*k*_−_ = 0).

**Fig. S3.**
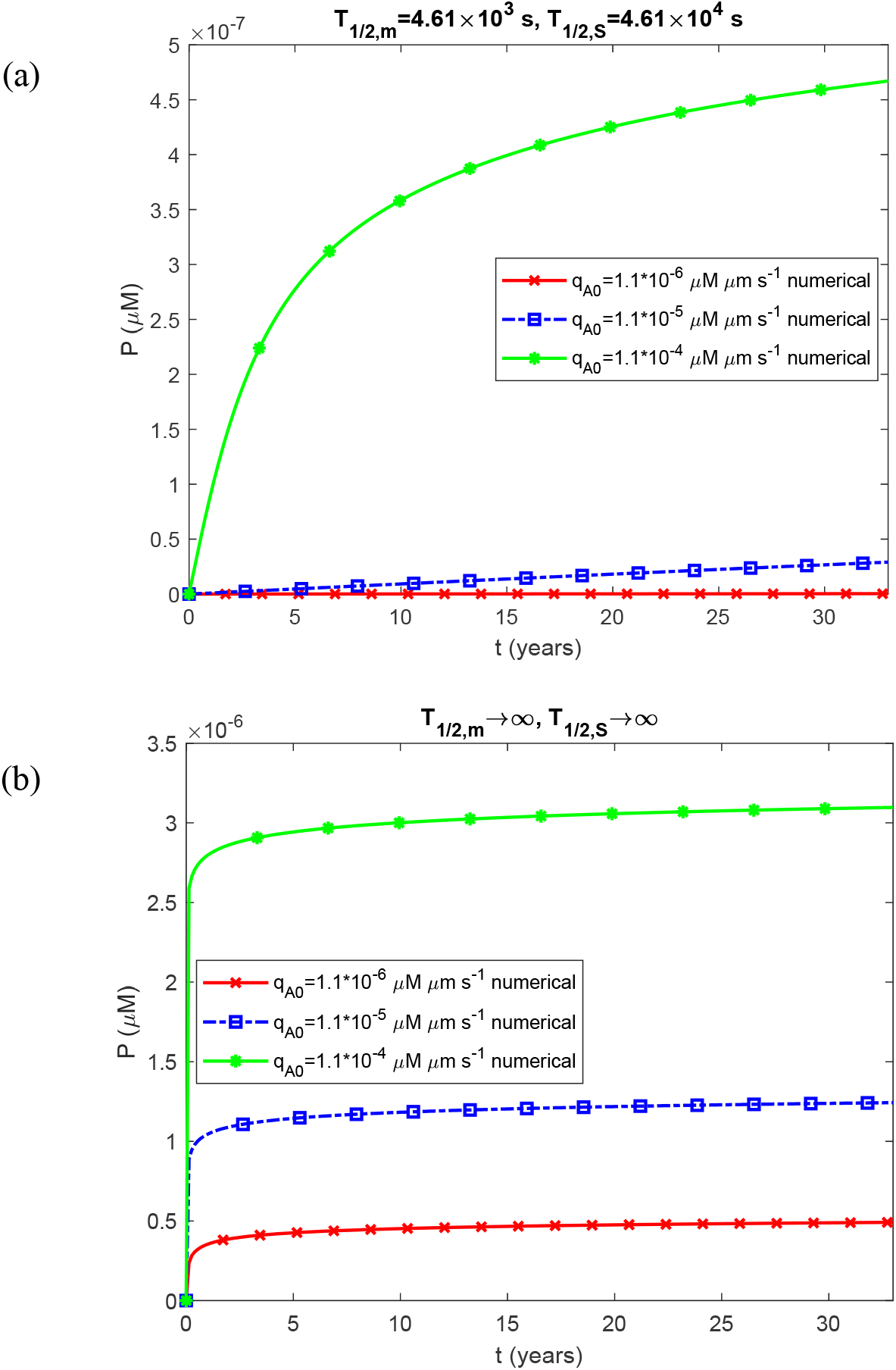
Molar concentration of Aβ fibrillar species of varying length as a function of time, *P*(*t*). The case in which oligomer dissociation into monomers is accounted for (*k*_*d*1_ = 3×10^−6^ s^-1^, *k*_*d*2_ = 1.94 ×10^−5^ μM^-1^ s^-1^) and fibril fragmentation is neglected (*k*_−_ = 0).

**Fig. S4.**
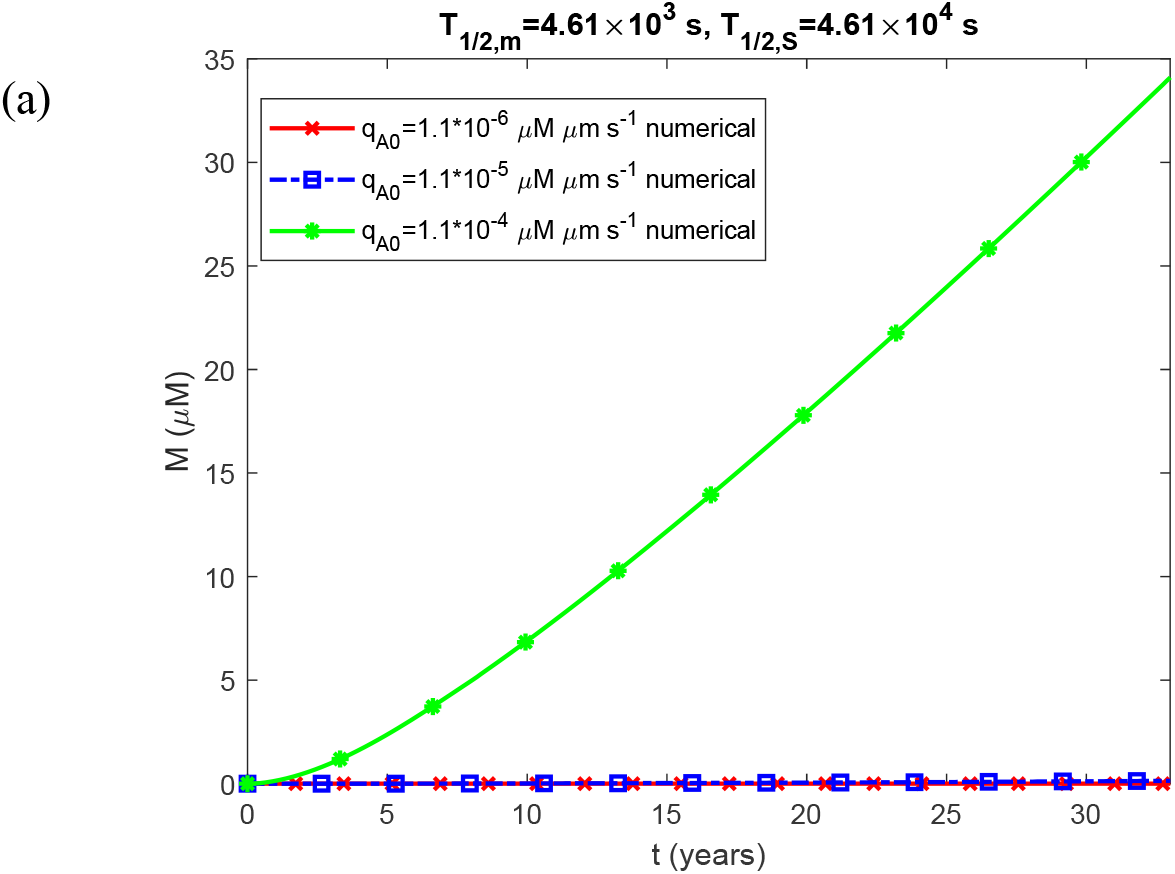
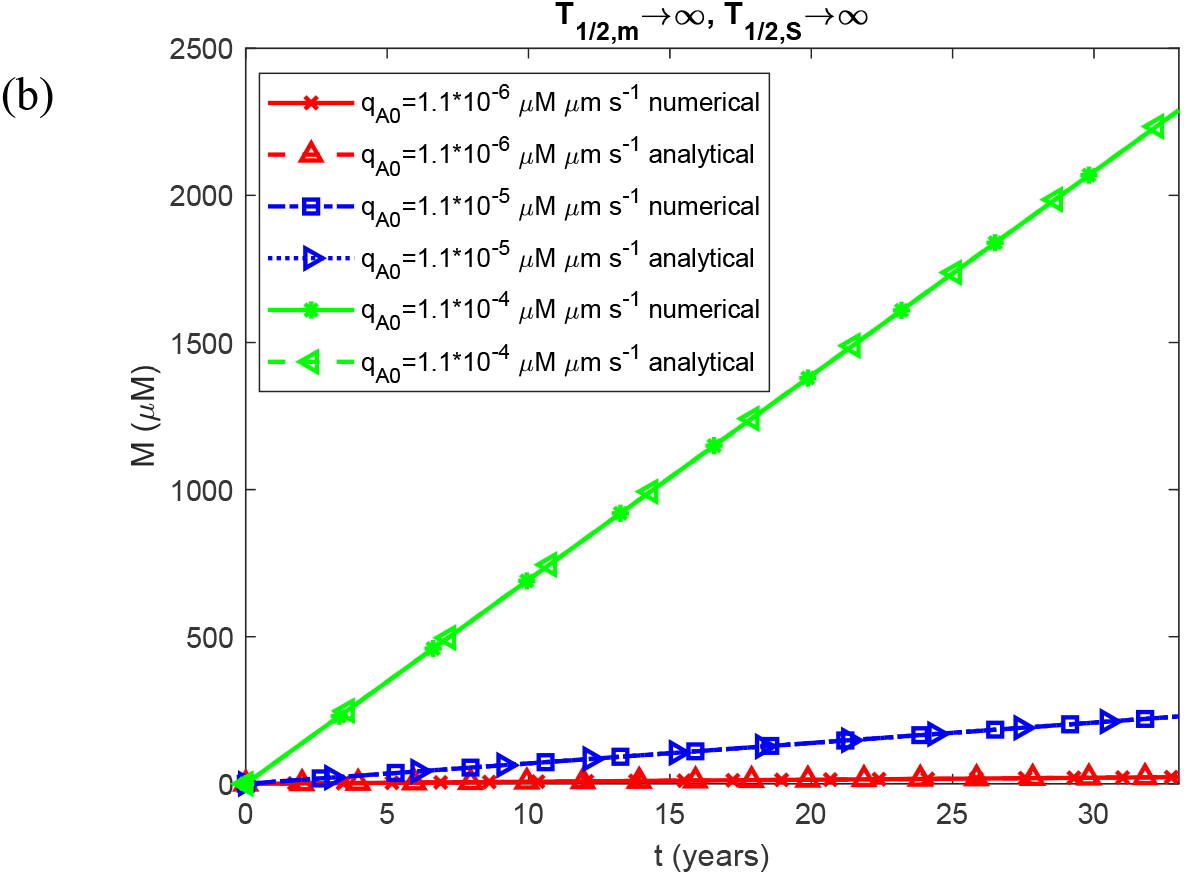
Molar concentration of Aβ monomers incorporated into fibrillar species of varying length as a function of time, *M* (*t*). The case in which oligomer dissociation into monomers is accounted for (*k*_*d*1_ = 3×10^−6^ s^-1^, *k*_*d*2_ = 1.94 ×10^−5^ μM^-1^ s^-1^) and fibril fragmentation is neglected (*k*_−_ = 0).

**Fig. S5.**
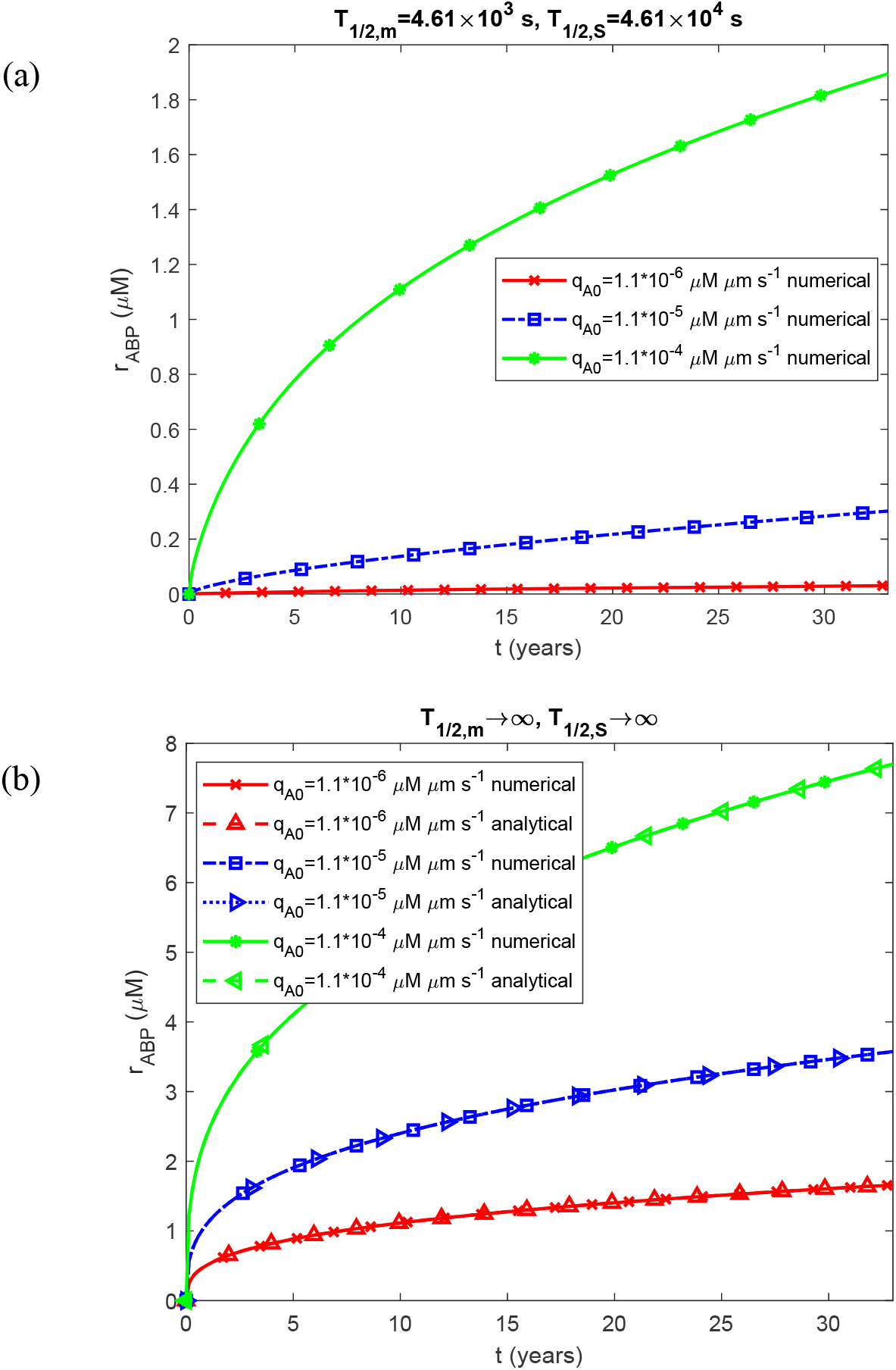
Radius of the Aβ plaque as a function of time, *r*_*ABP*_ (*t*). The case in which oligomer dissociation into monomers is accounted for (*k*_*d*1_ = 3×10^−6^ s^-1^, *k*_*d*2_ = 1.94 ×10^−5^ μM^-1^ s^-1^) and fibril fragmentation is neglected (*k*_−_ = 0).

**Fig. S6.**
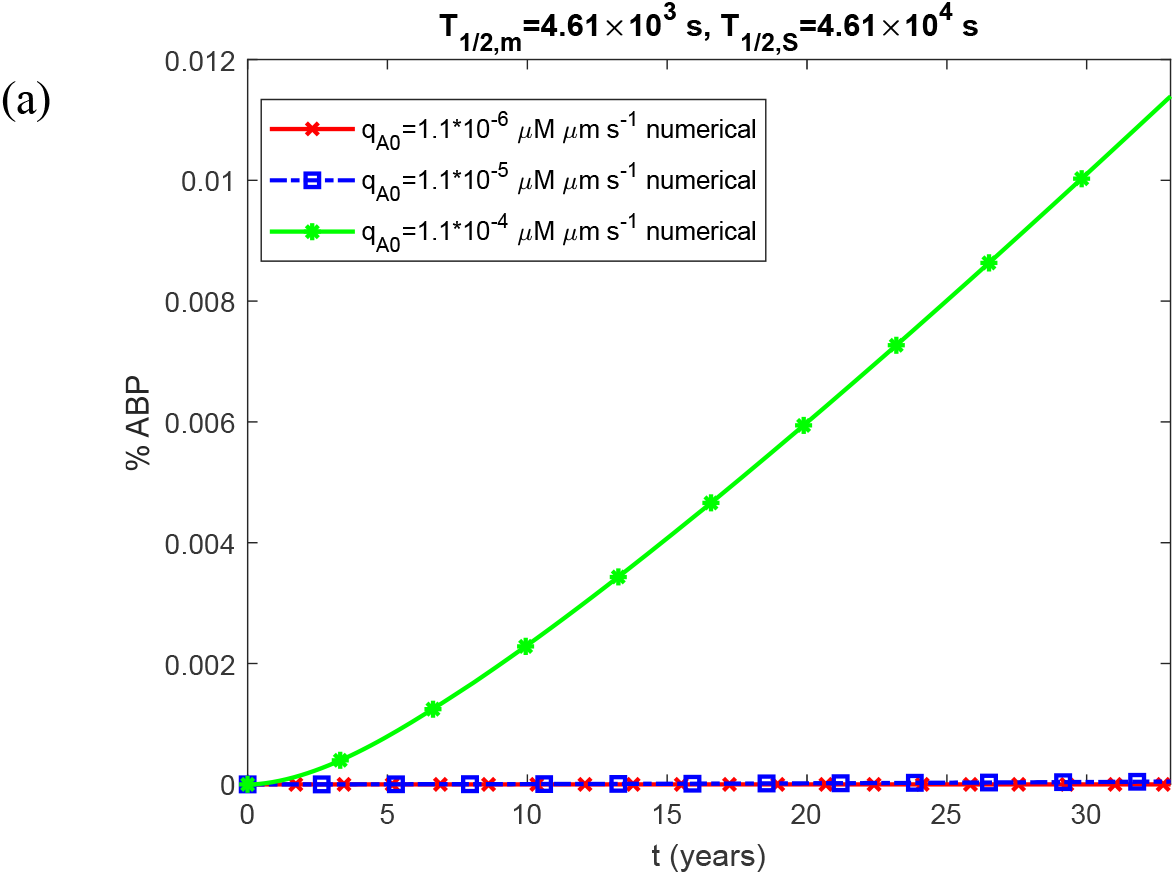
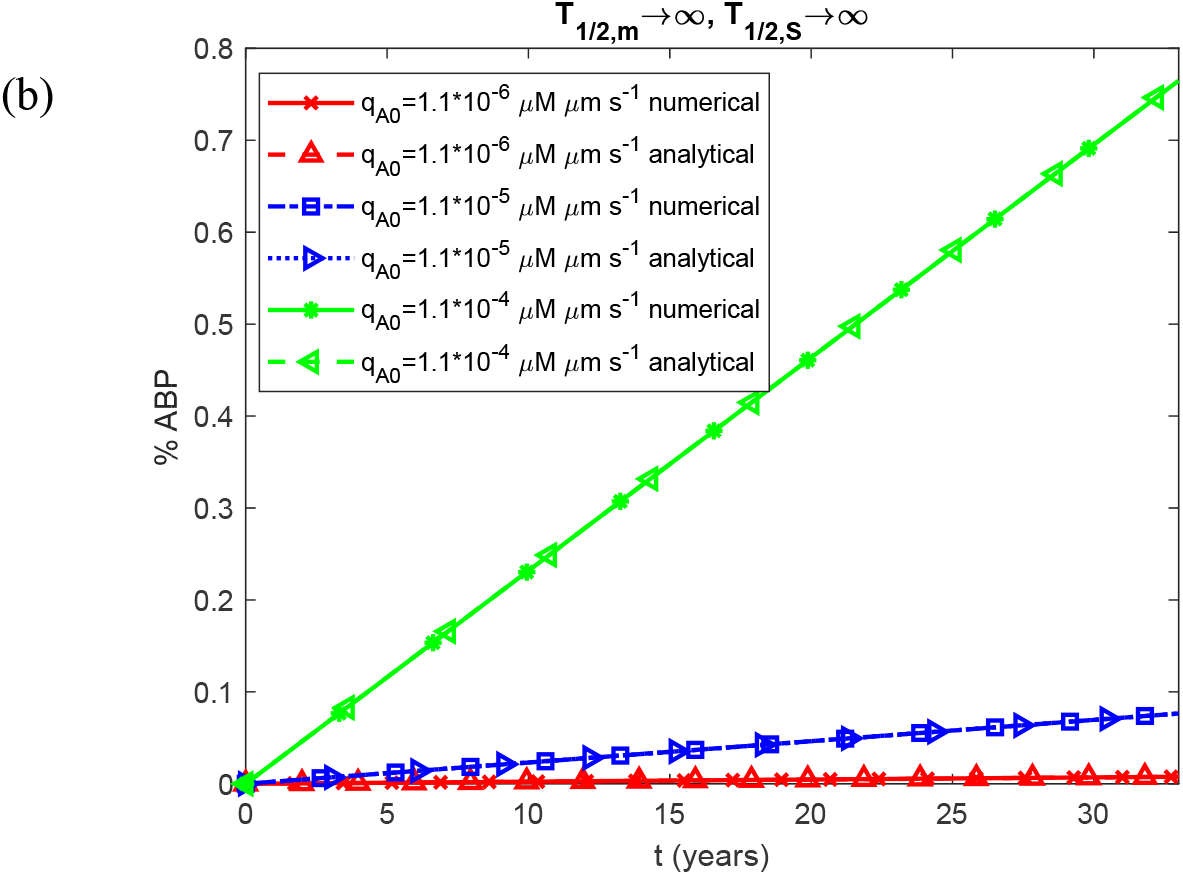
Percentage of the CV occupied by the Aβ plaque as a function of time. The case in which oligomer dissociation into monomers is accounted for (*k*_*d*1_ = 3×10^−6^ s^-1^, *k*_*d*2_ = 1.94 ×10^−5^ μM^-1^ s^-1^) and fibril fragmentation is neglected (*k*_−_ = 0).

**Fig. S7.**
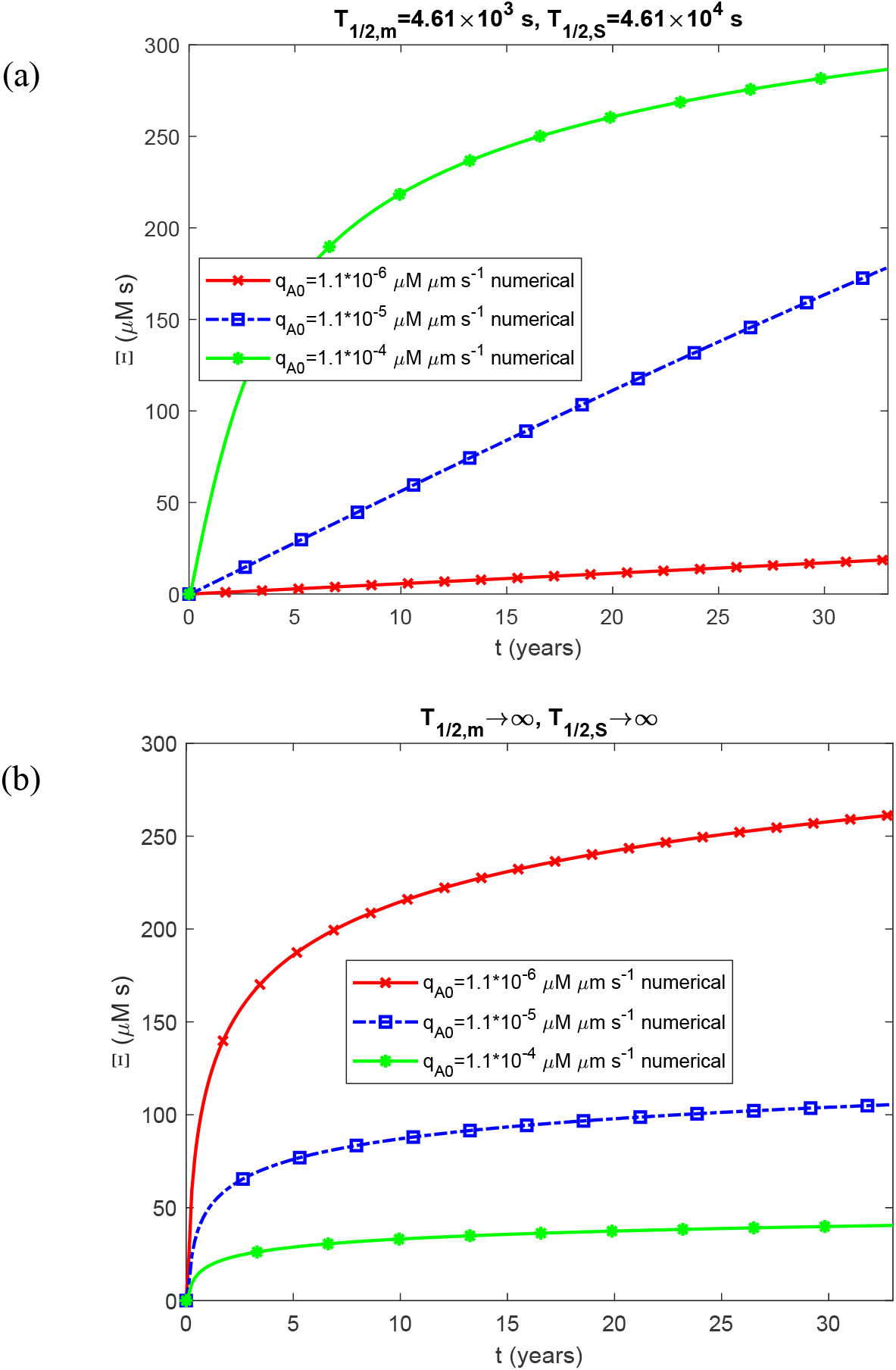
Accumulated neurotoxicity as a function of time, Ξ(*t*). The case in which oligomer dissociation into monomers is accounted for (*k*_*d*1_ = 3×10^−6^ s^-1^, *k*_*d*2_ = 1.94 ×10^−5^ μM^-1^ s^-1^) and fibril fragmentation is neglected (*k*_−_ = 0).

**Fig. S8.**
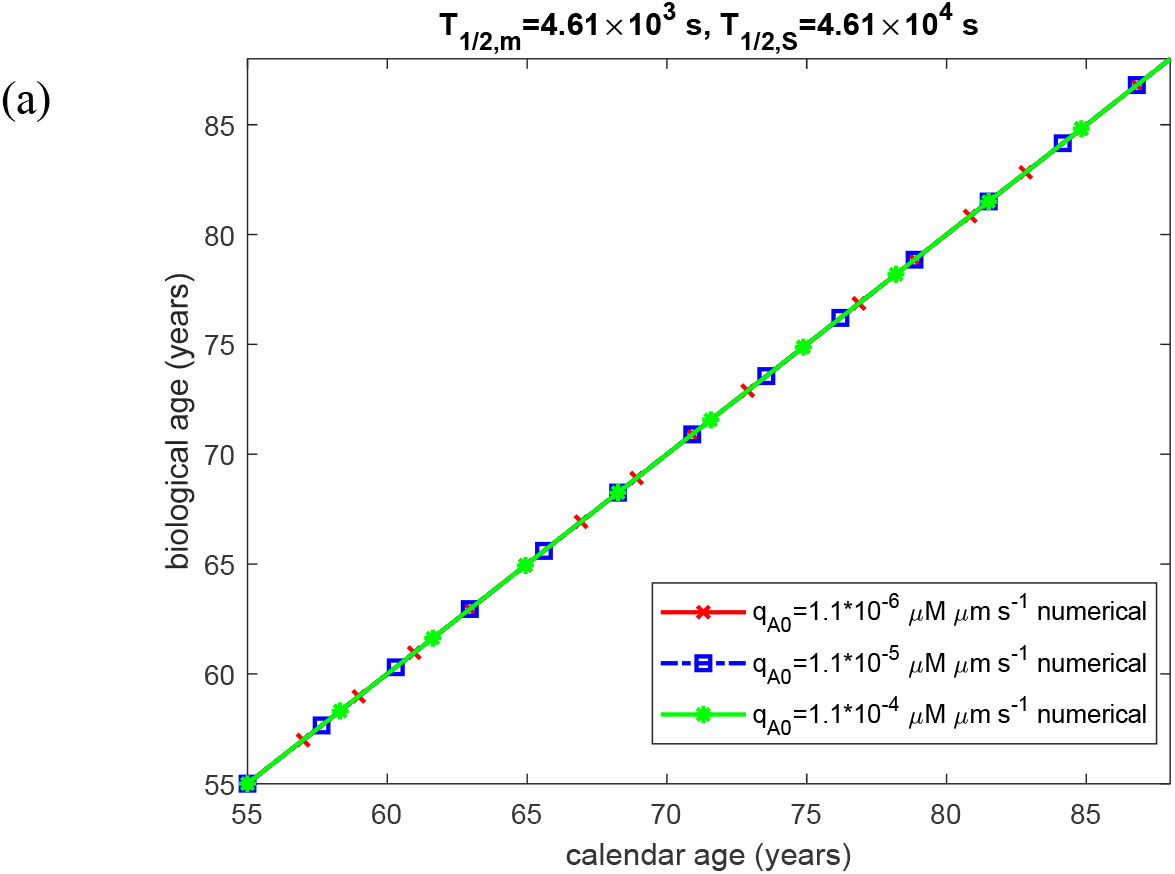
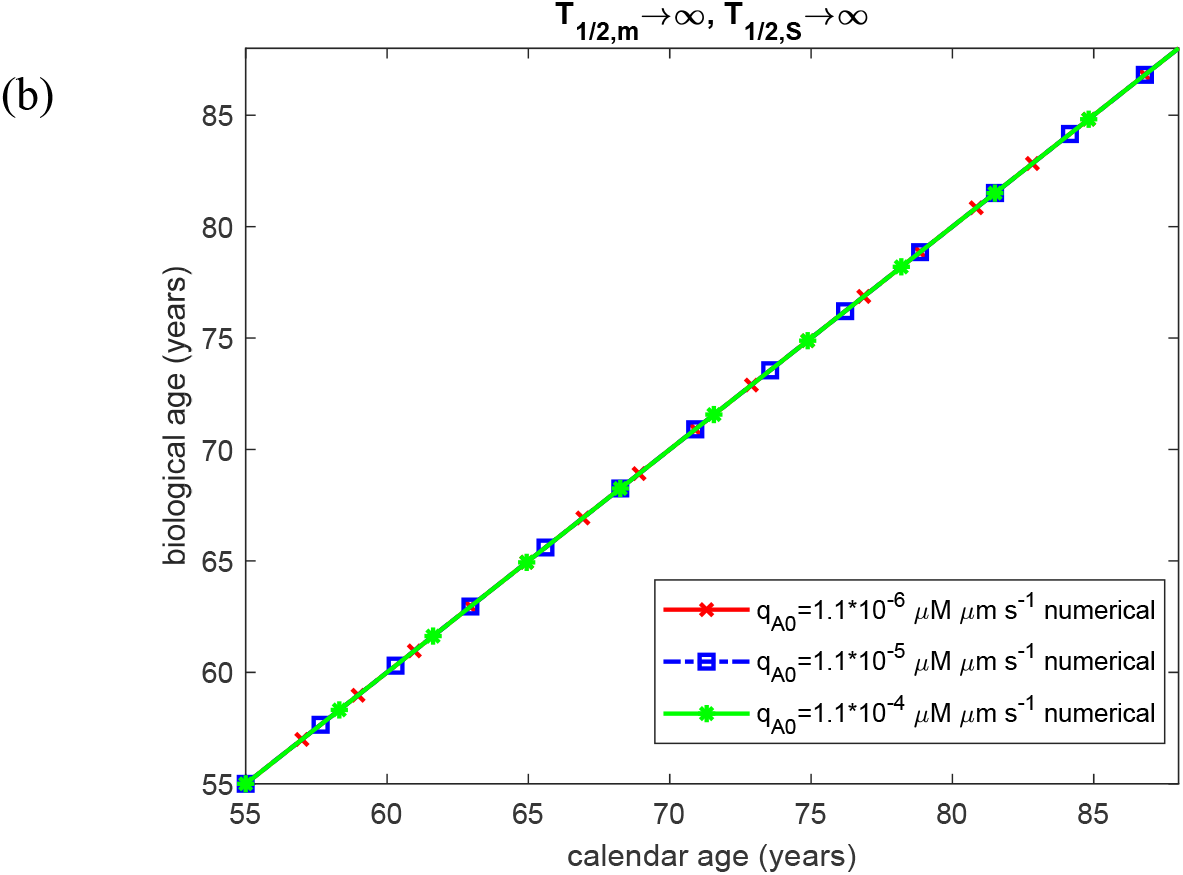
Biological age as a function of calendar age. The case in which oligomer dissociation into monomers is accounted for (*k*_*d*1_ = 3×10^−6^ s^-1^, *k*_*d*2_ = 1.94 ×10^−5^ μM^-1^ s^-1^) and fibril fragmentation is neglected (*k*_−_ = 0).

**Fig. S9.**
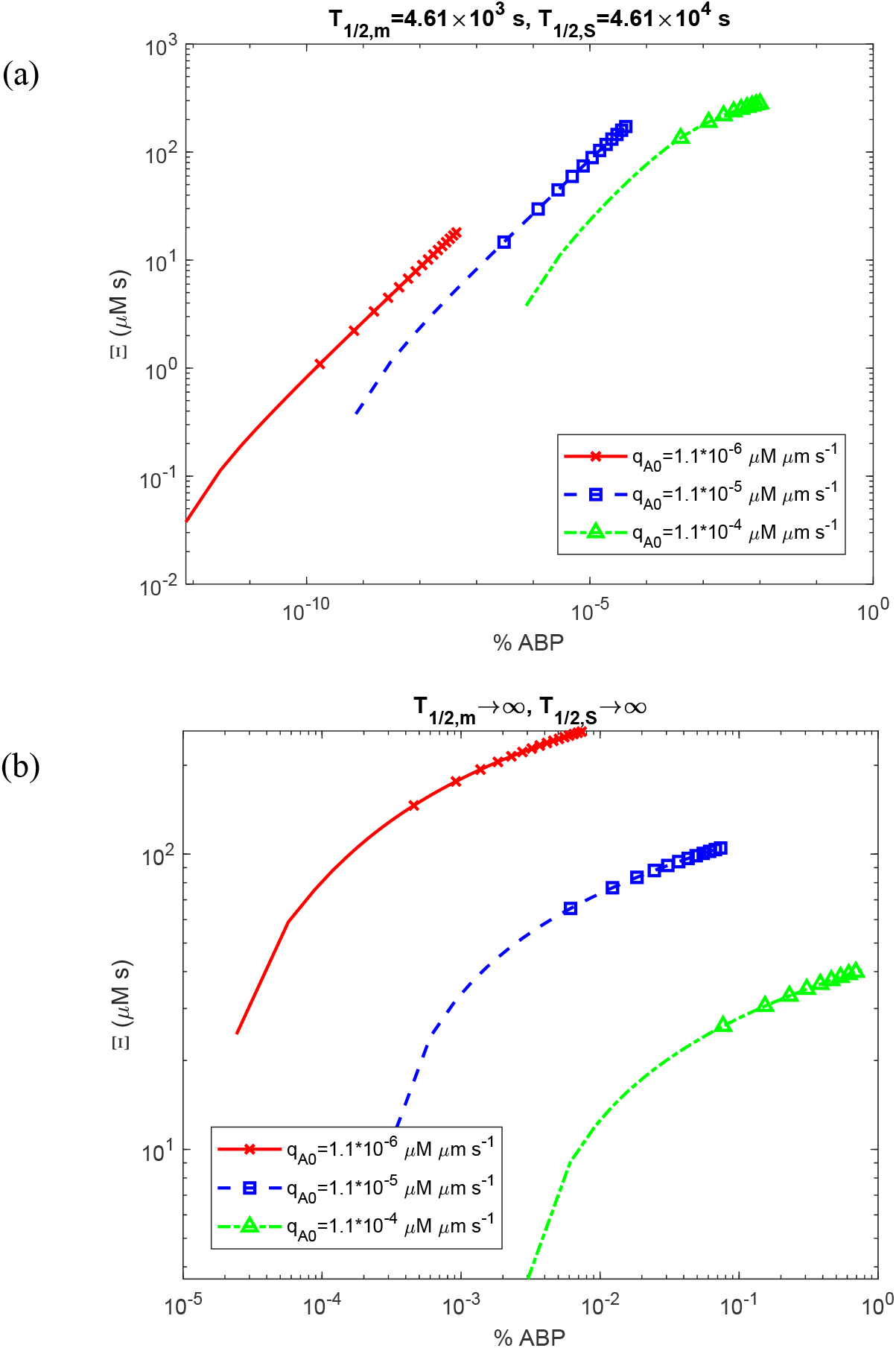
Accumulated neurotoxicity as a function of the percentage of the CV occupied by the Aβ plaque. The case in which oligomer dissociation into monomers is accounted for (*k*_*d*1_ = 3×10^−6^ s^-1^, *k*_*d*2_ = 1.94 ×10^−5^ μM^-1^ s^-1^) and fibril fragmentation is neglected (*k*_−_ = 0).

**Fig. S10.**
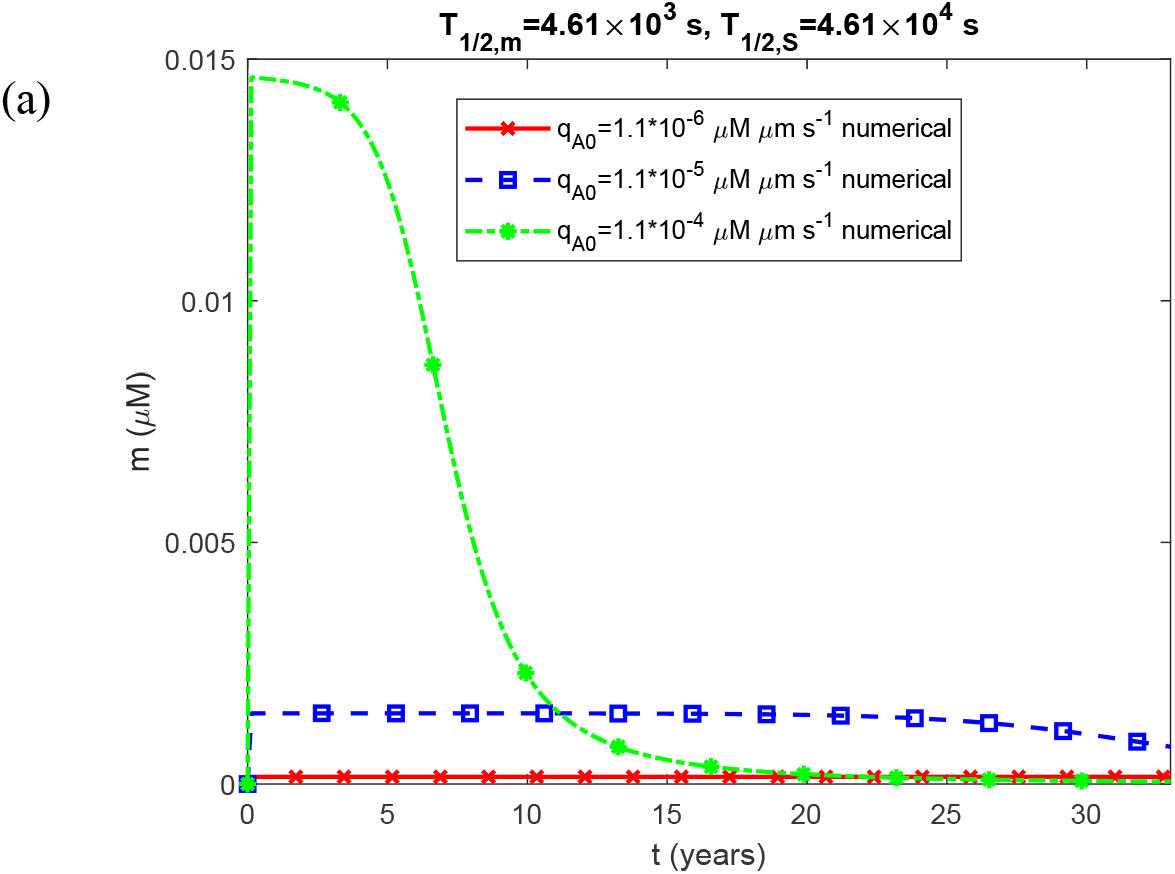
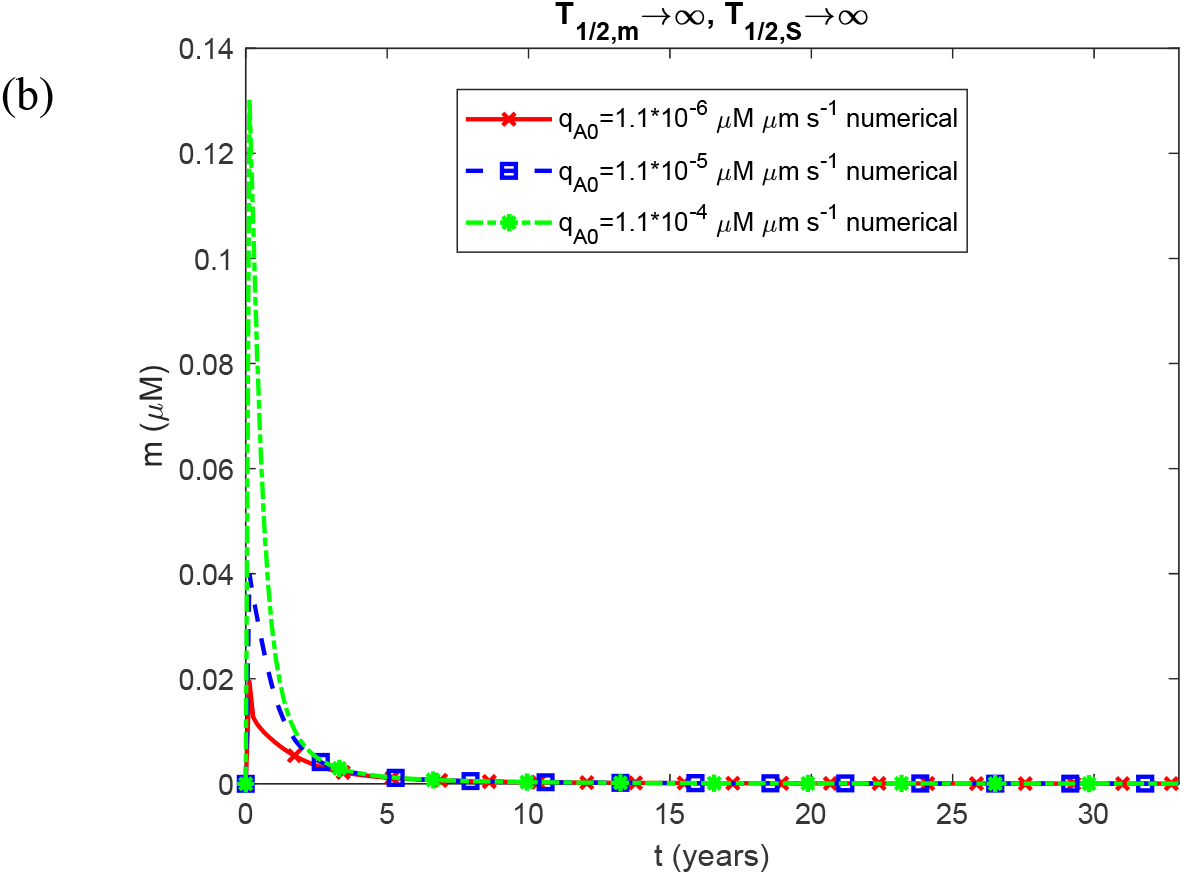
Molar concentration of free Aβ monomers as a function of time, *m*(*t*). The case in which both oligomer dissociation into monomers (*k*_*d*1_ = 3×10^−6^ s^-1^, *k*_*d*2_ = 1.94 ×10^−5^ μM^-1^ s^-1^) and fibril fragmentation (*k*_−_ = 10^−14^ s^-1^) are accounted for.

**Fig. S11.**
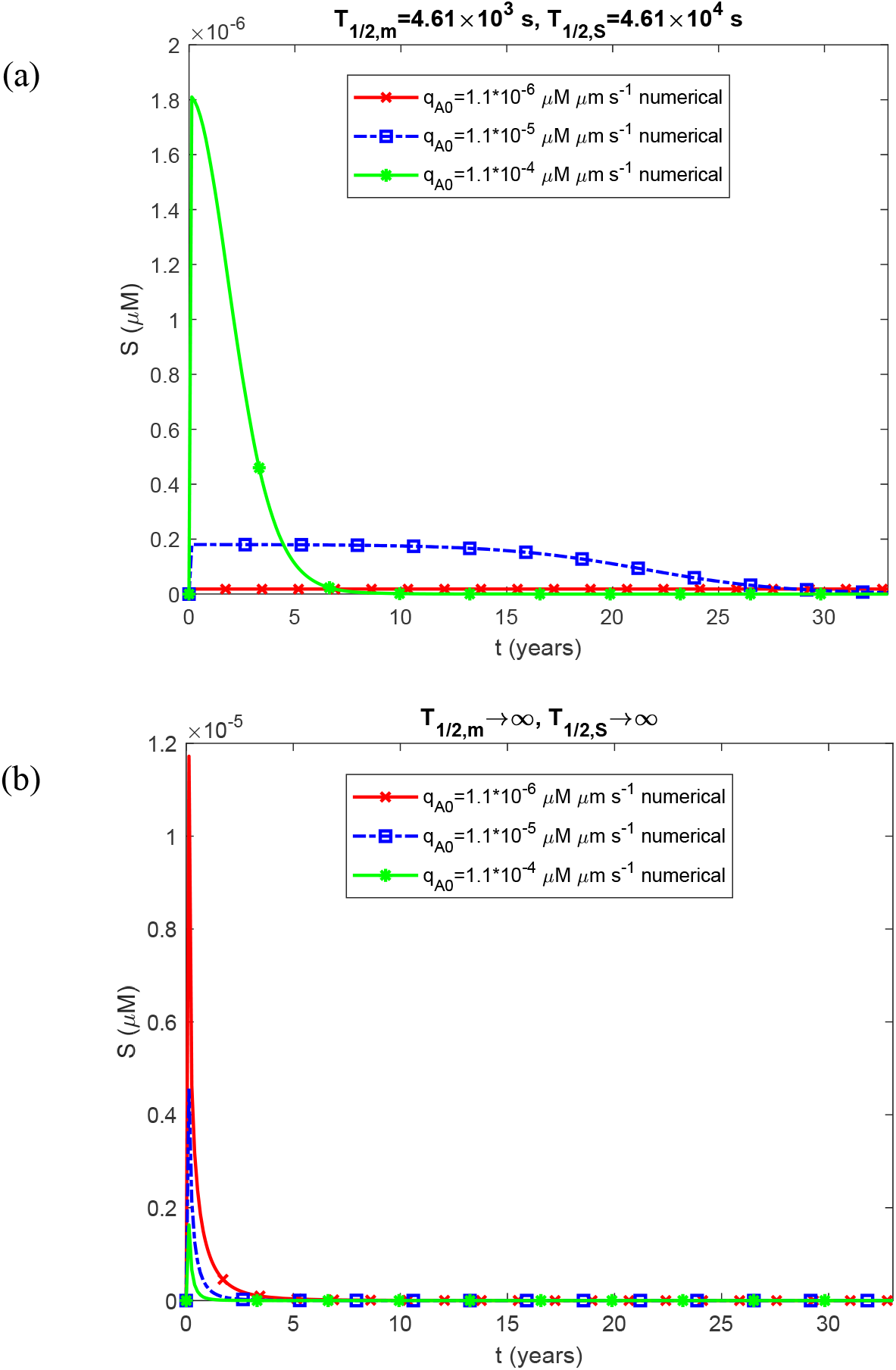
Molar concentration of free on-pathway Aβ oligomers as a function of time, *S* (*t*). The case in which both oligomer dissociation into monomers (*k* _*d*1_= 3×10^−6^ s^-1^, *k*_*d*2_ = 1.94 ×10^−5^ μM^-1^ s^-1^) and fibril fragmentation (*k*_−_ = 10^−14^ s^-1^) are accounted for.

**Fig. S12.**
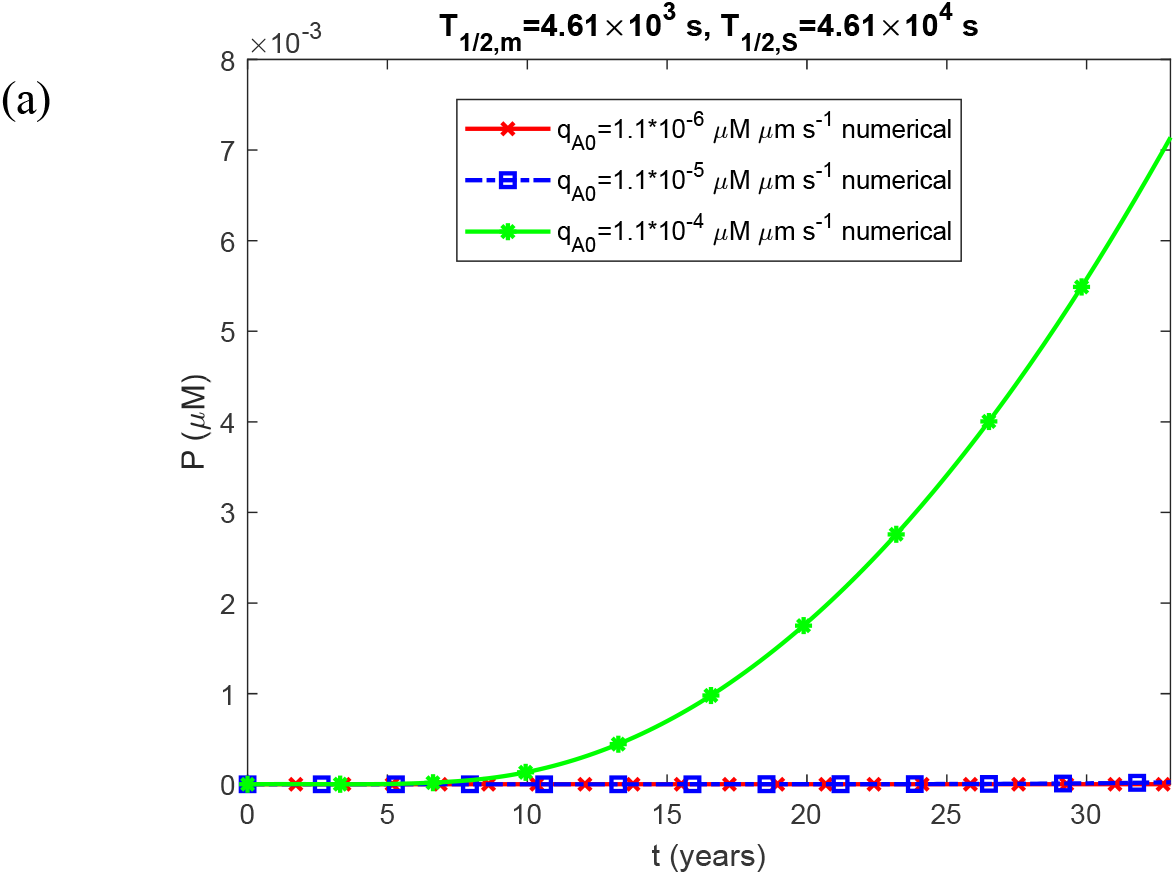
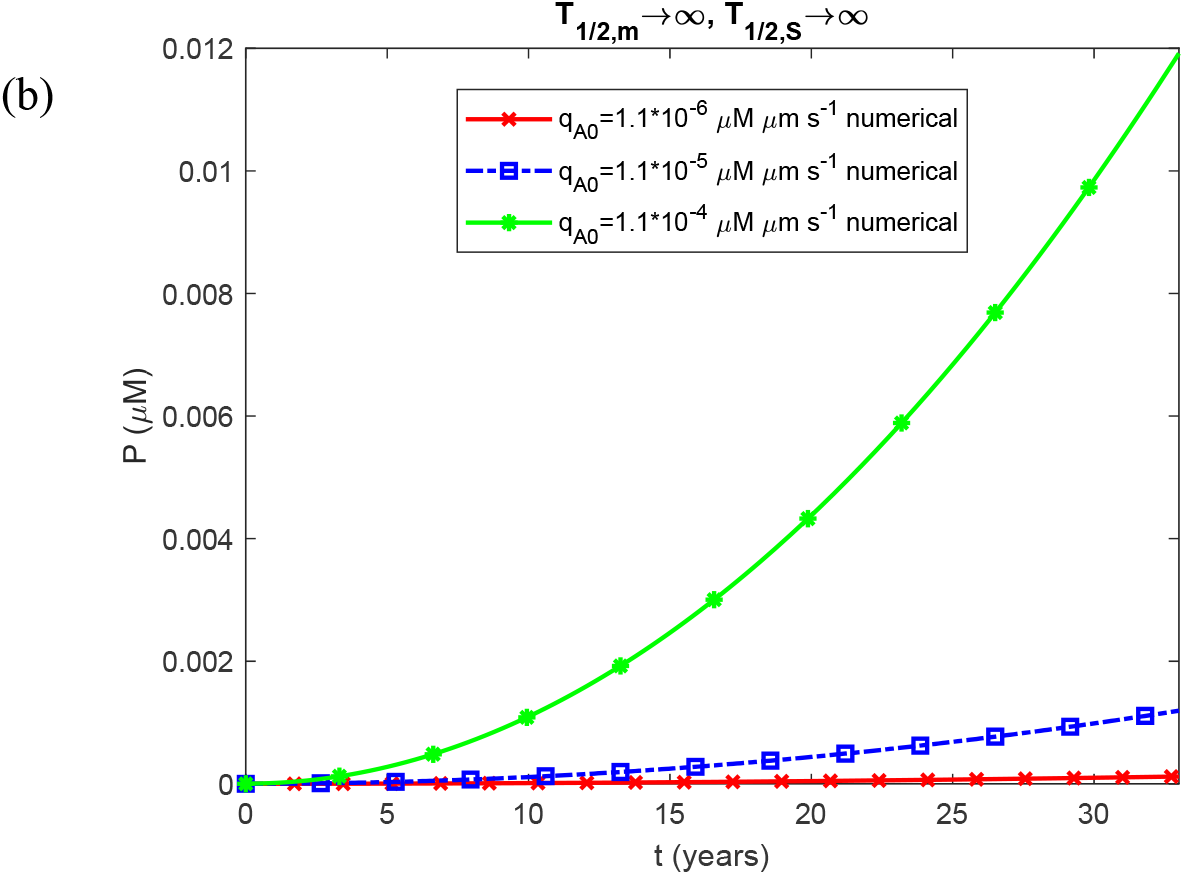
Molar concentration of Aβ fibrillar species of varying length as a function of time, *P*(*t*). The case in which both oligomer dissociation into monomers (*k* _*d*1_= 3×10^−6^ s^-1^, *k*_*d*2_ = 1.94 ×10^−5^ μM^-1^ s^-1^) and fibril fragmentation (*k*_−_ = 10^−14^ s^-1^) are accounted for.

**Fig. S13.**
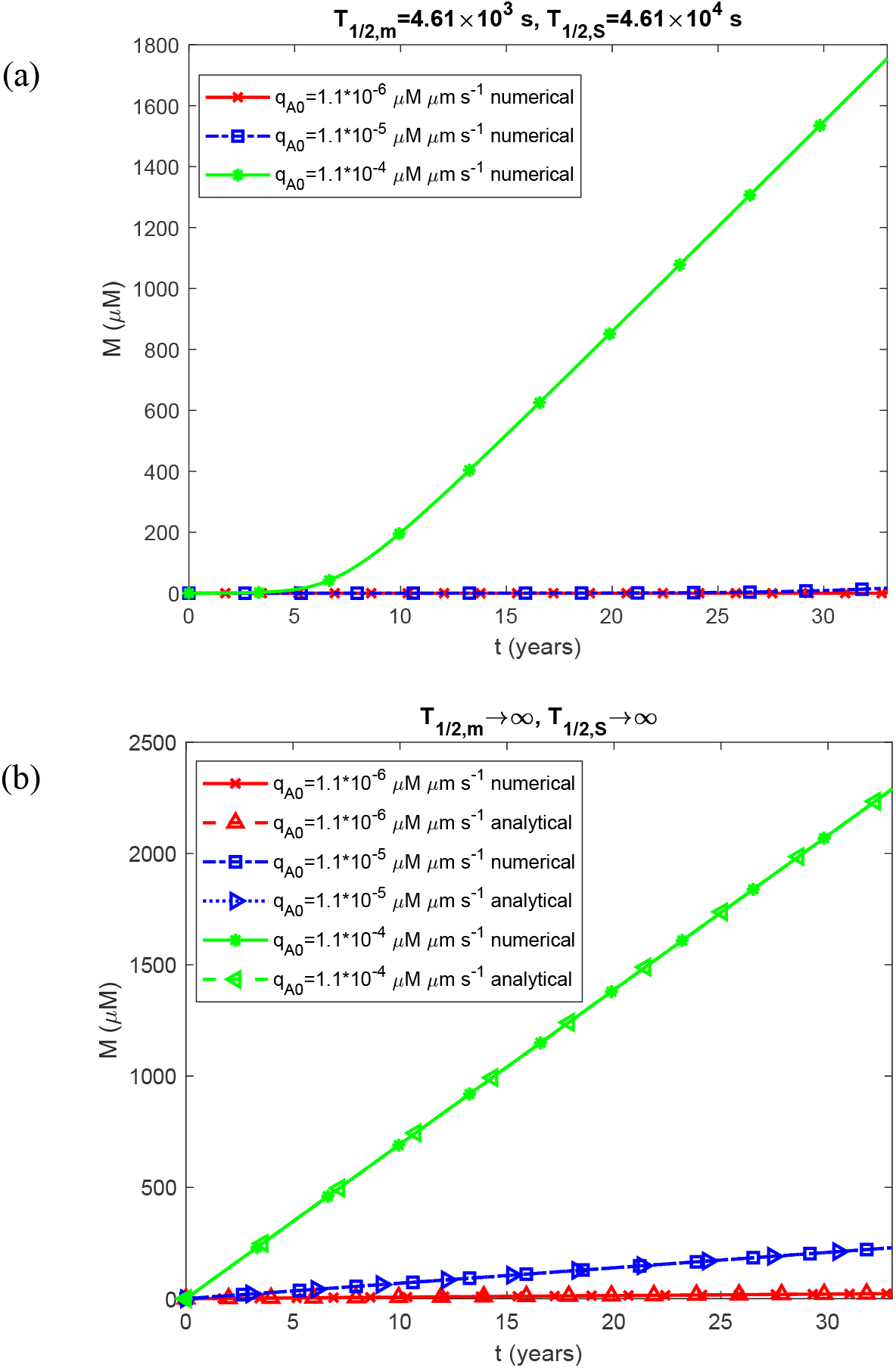
Molar concentration of Aβ monomers incorporated into fibrillar species of varying length as a function of time, *M* (*t*). The case in which both oligomer dissociation into monomers (*k*_*d*1_ = 3×10−6 s^-1^, *k*_*d*12_ = 1.94 ×10^−5^ μM^-1^ s^-1^) and fibril fragmentation (*k*_−_ = 10^−14^ s^-1^) are accounted for.

**Fig. S14.**
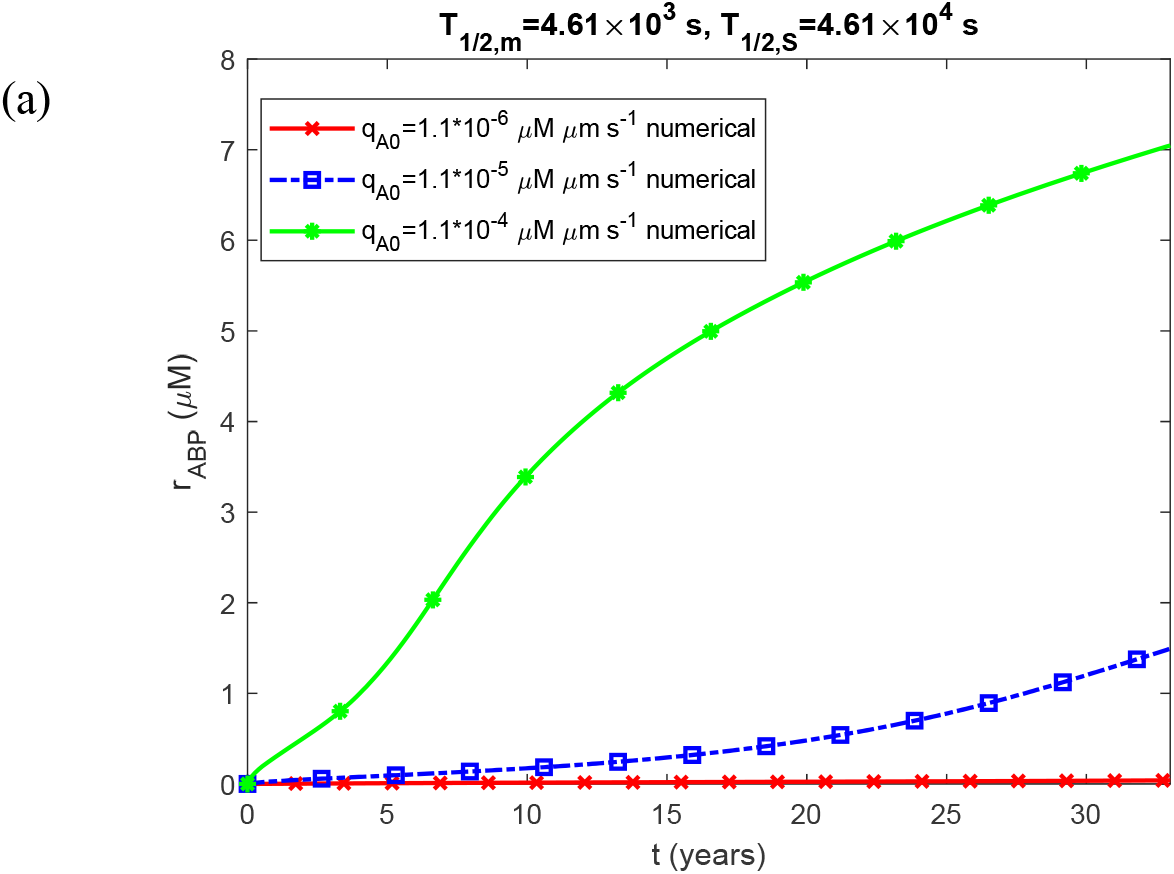
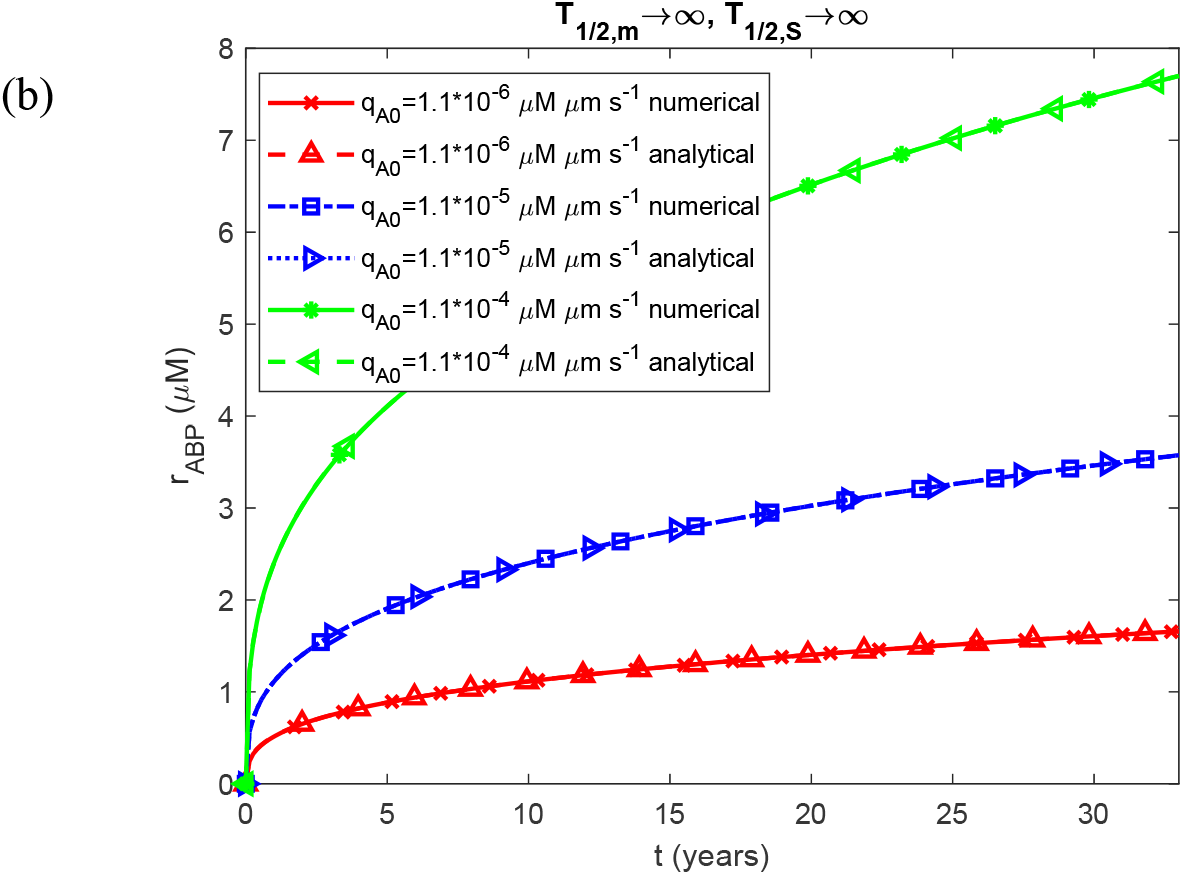
Radius of the Aβ plaque as a function of time, *r*_*ABP*_ (*t*). The case in which both oligomer dissociation into monomers (*k*_*d*1_ = 3×10^−6^ s^-1^, *k*_*d*2_ = 1.94 ×10^−5^ μM^-1^ s^-1^) and fibril fragmentation (*k*_−_ = 10^−14^ s^-1^) are accounted for.

**Fig. S15.**
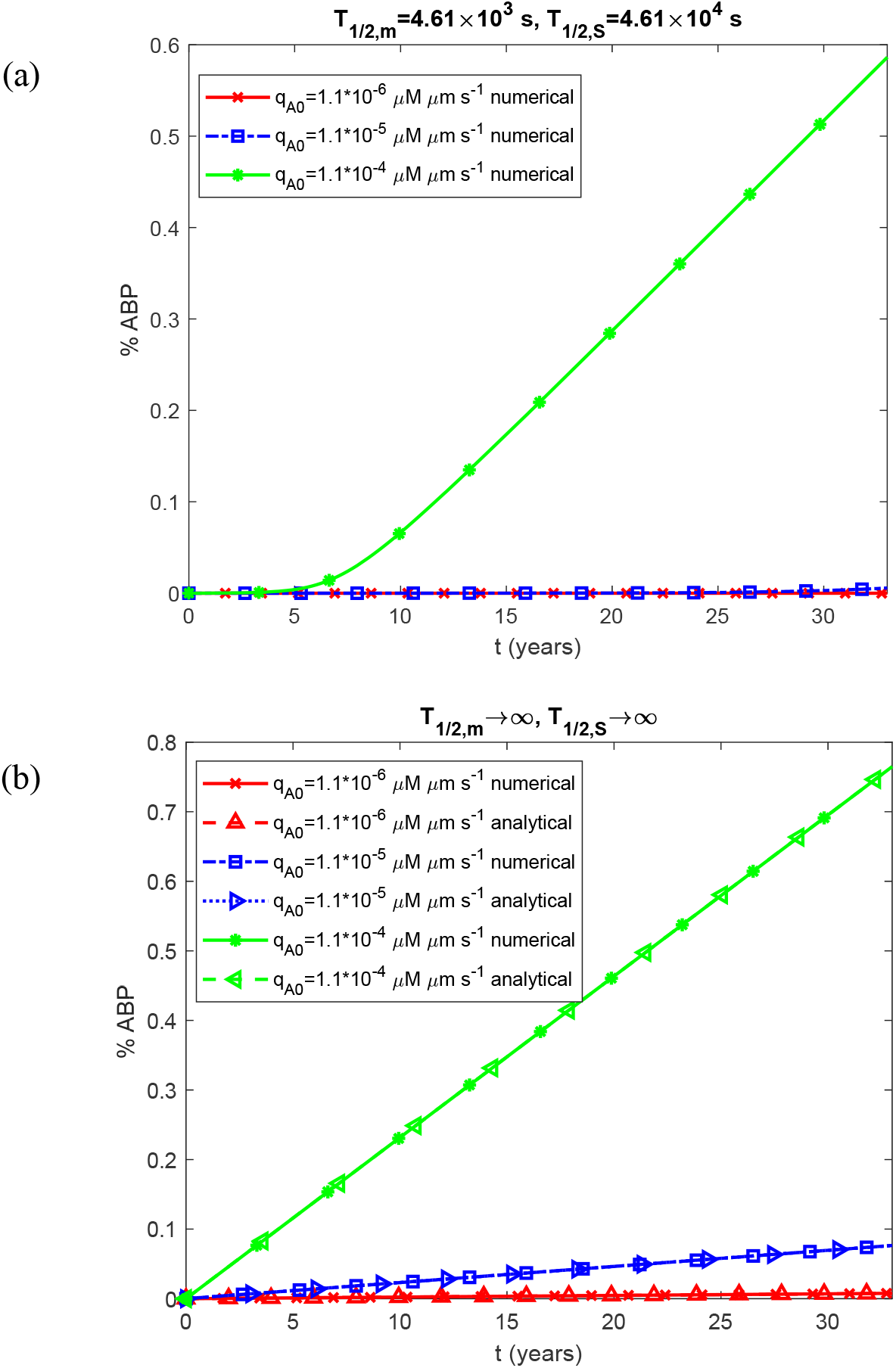
Percentage of the CV occupied by the Aβ plaque as a function of time. The case in which both oligomer dissociation into monomers (*k*_*d*1_ = 3×10^−6^ s^-1^, *k*_*d*2_ = 1.94 ×10^−5^ μM^-1^ s^-1^) and fibril fragmentation (*k*_−_ = 10^−14^ s^-1^) are accounted for.

**Fig. S16.**
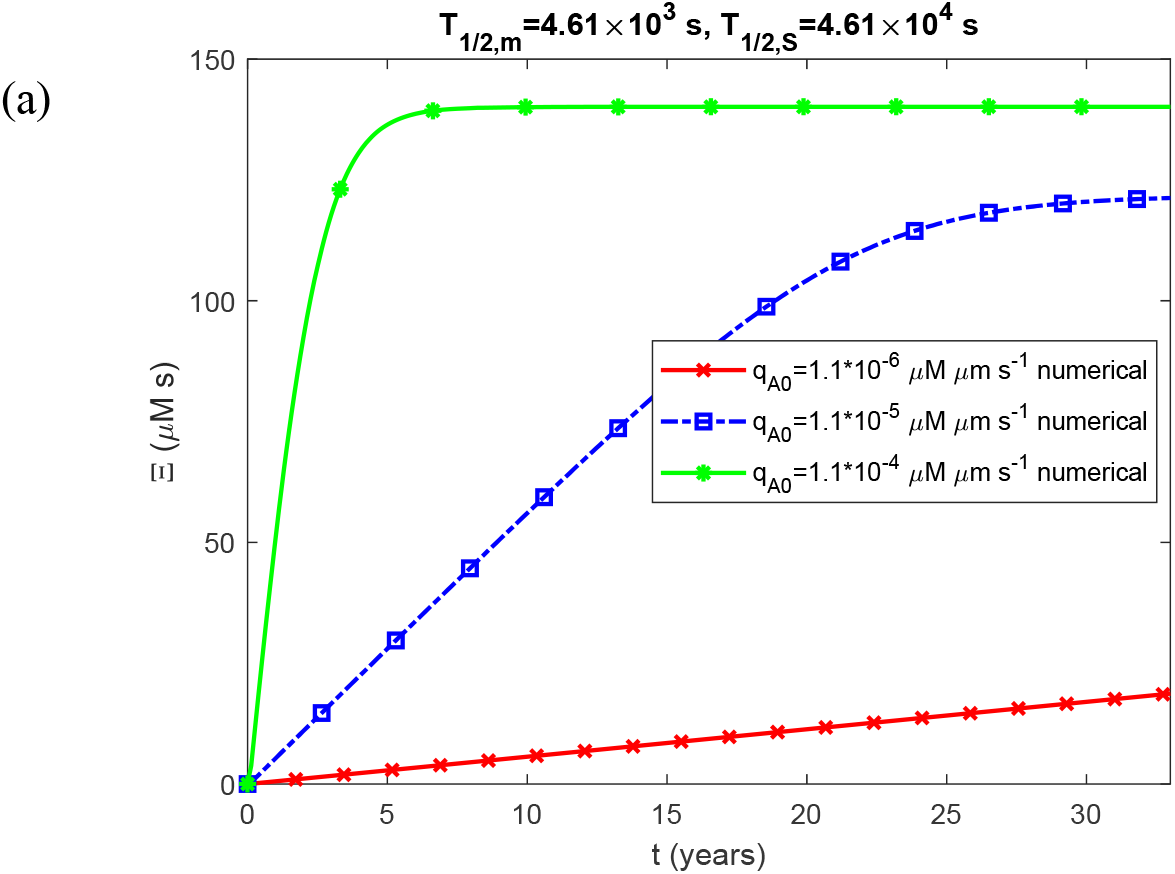
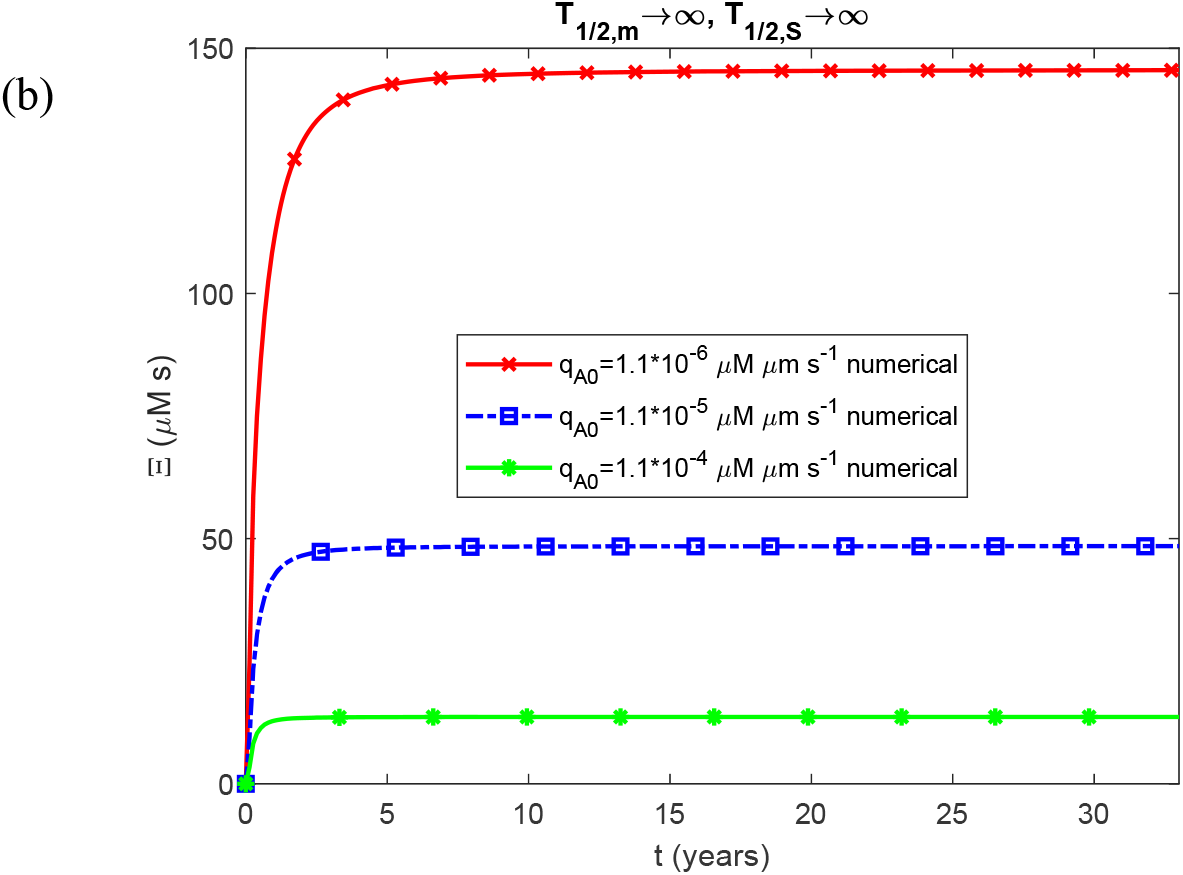
Accumulated neurotoxicity as a function of time, Ξ(*t*). The case in which both oligomer dissociation into monomers (*k*_*d*1_ = 3×10^−6^ s^-1^, *k*_*d*2_ = 1.94 ×10^−5^ μM^-1^ s^-1^) and fibril fragmentation (*k*_−_ = 10^−14^ s^-1^) are accounted for.

**Fig. S17.**
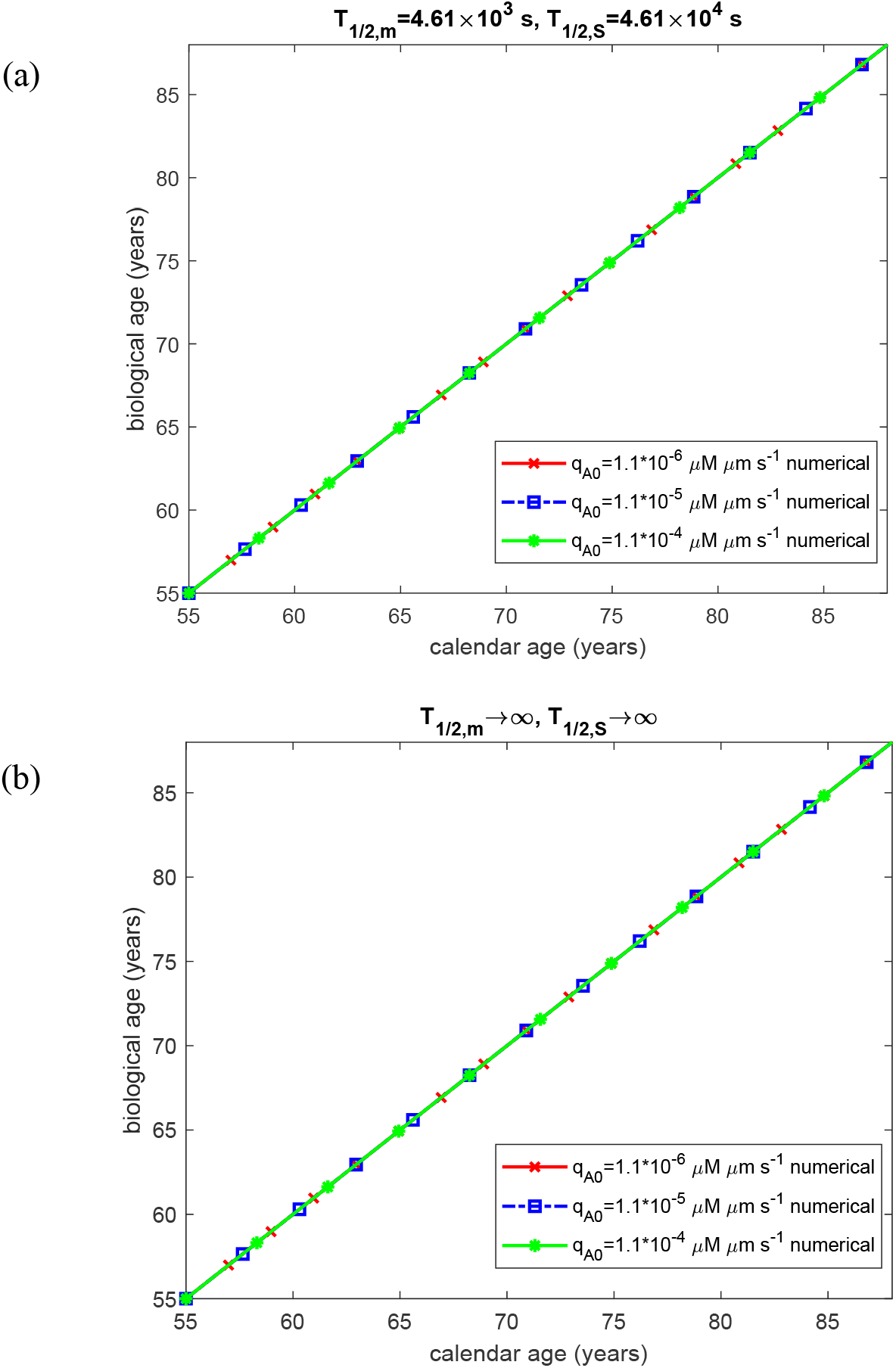
Biological age as a function of calendar age. The case in which both oligomer dissociation into monomers (*k*_*d*1_ = 3×10^−6^ s^-1^, *k*_*d*2_ = 1.94 ×10^−5^ μM^-1^ s^-1^) and fibril fragmentation (*k*_−_ = 10^−14^ s^-1^) are accounted for.

**Fig. S18.**
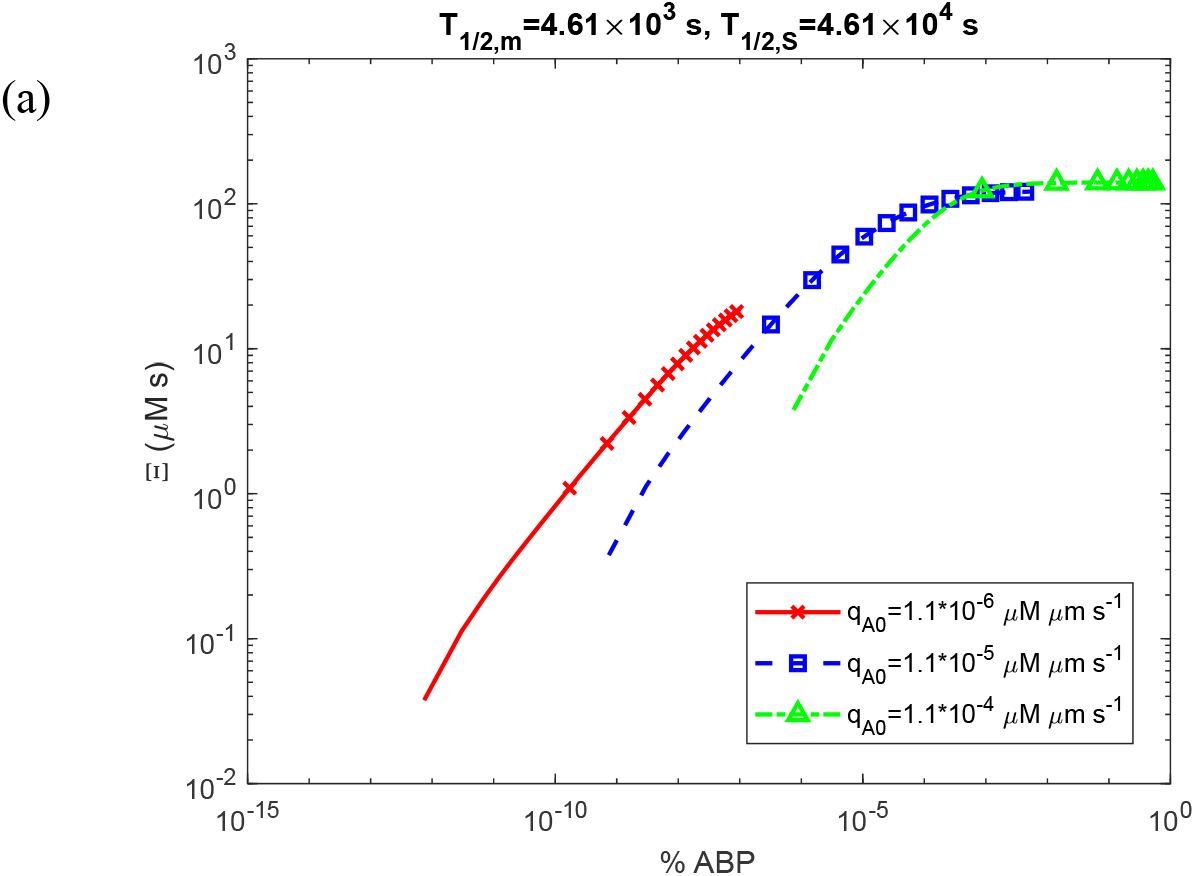
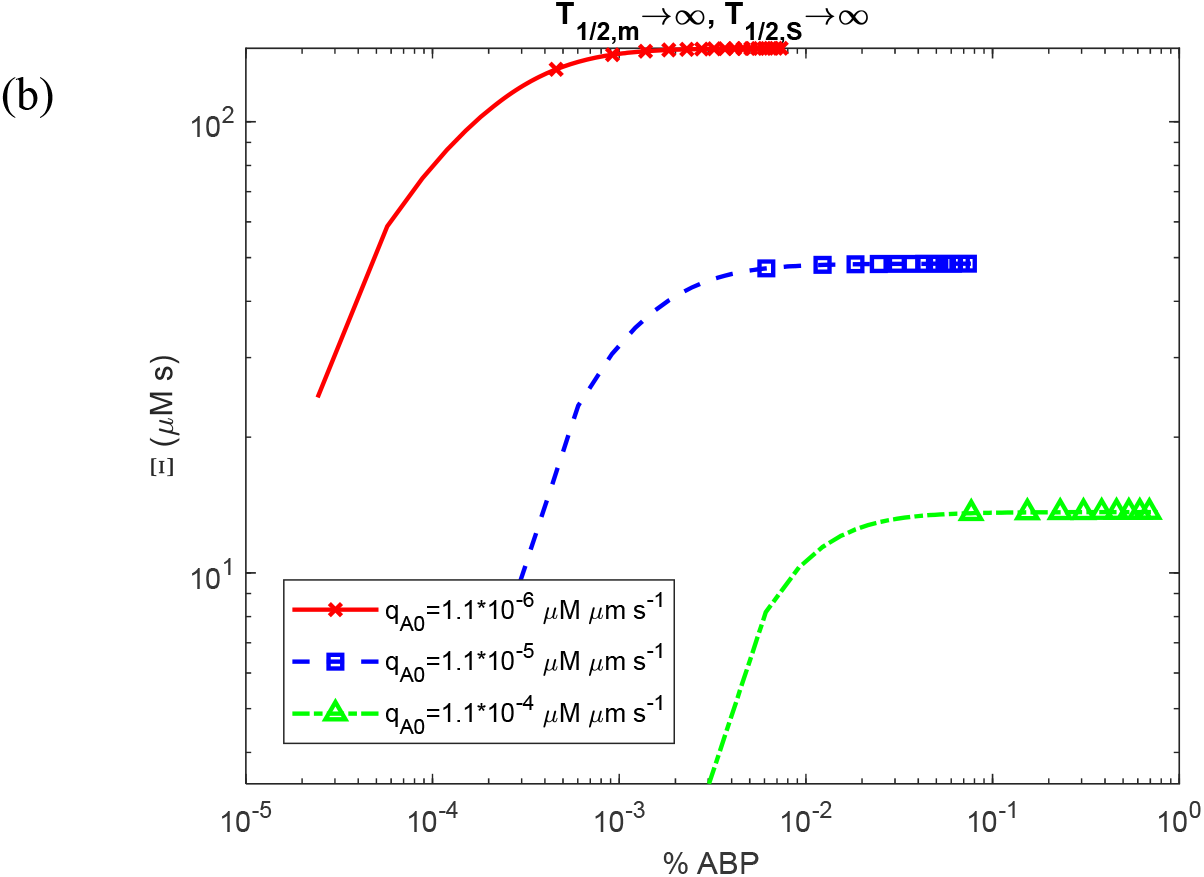
Accumulated neurotoxicity as a function of the percentage of the CV occupied by the Aβ plaque. The case in which both oligomer dissociation into monomers (*k*_*d*1_ = 3×10^−6^ s^-1^, *k*_*d*2_ = 1.94 ×10^−5^ μM^-1^ s^-1^) and fibril fragmentation (*k*_−_ = 10^−14^ s^-1^) are accounted for.

